# Multiomic definition of generalizable endotypes in human acute pancreatitis

**DOI:** 10.1101/539569

**Authors:** Lucile P. A. Neyton, Xiaozhong Zheng, Christos Skouras, Andrea Doeschl-Wilson, Michael U. Gutmann, Christopher Yau, Iain Uings, Francesco V. Rao, Armel Nicolas, Craig Marshall, Lisa-Marie Wilson, J. Kenneth Baillie, Damian J. Mole

## Abstract

Acute pancreatitis (AP) is sudden onset pancreas inflammation that causes multiple organ dysfunction syndrome (MODS) and death in certain individuals who develop AP yet minimal systemic inflammation in others. Here, we show that this observed diversity in systemic response and outcome is accompagnied by diversity in molecular subtypes that can be identified using computational analysis of clinical and multiomic data. We integrated co-incident whole blood transcriptomic, plasma proteomic, and serum metabolomic data at serial time points from a cohort of patients presenting with AP and systematically evaluated four different metrics for patient similarity, using unbiased mathematical, biological and clinical measures of internal and external validity. Our results identify four distinct and stable AP endotypes that are characterized by pathway and biomarker combination stereotypes into hypermetabolic, hepatopancreaticobiliary, catabolic and innate immune endotypes. The catabolic endotype in AP strikingly recapitulates a disease endotype previously reported in acute respiratory distress syndrome, a recognized complication of AP. Our findings demonstrate that clinically-relevant and generalizable endotypes exist in AP.

## Main

Acute pancreatitis (AP) is defined as acute inflammation of the pancreas gland^1^. AP has a worldwide incidence of 34 per 100,000 person-years^2^, and is the commonest gastrointestinal cause for emergency hospital admission^3^. Inflammatory damage to pancreatic acinar cells initiates an inflammatory cascade mediated by damage-associated molecular patterns, alarmins, inflammatory cytokines, metabolites and other soluble and cellular mediators of inflammation that propagate inflammation locally in the pancreas, and cause extrapancreatic organ dysfunction in the lungs, kidney and liver and other body systems, together resulting in multiple organ dysfunction syndrome (MODS)^4,5^. MODS occurs in 1 in 4 individuals who develop AP and is accompanied by deregulation of cardiovascular, autonomic nervous and immune system homeostasis^6^, leading to death in one fifth of those with AP-MODS^7^. Despite this currently accepted unifying disease model, each person who develops AP has a severity pattern of systemic inflammation and MODS that is unique and is not directly proportional to the amount of pancreas damage on radiological imaging^8,9^, and this individualized response determines the disease outcome. This personalized response to AP is further nuanced by the diversity of etiologies and initiating events in AP, that include choledocholithiasis, excess ingestion of alcohol, trauma, pancreatic manipulation at endoscopy, viral infections, certain venoms and specific prescription medicines^6^. Currently, the clinicopathological paradigm in AP is convergent, whereby diverse etiologies converge onto acinar cell damage, and the resulting systemic inflammatory response is stratified as mild, moderate or severe (**Figure 1a**). We propose an alternative model that describes illness trajectories in AP, which if correct, will have greater clinical utility in guiding treatment. Specifically, we propose the existence of molecular subtypes in AP, designated as endotypes^10^, and hypothesize that detailed knowledge of those endotypes will have clinical and therapeutic relevance (**Figure 1b**). Importantly, different endotypes may present similar clinical features, and one phenotype may be shared by multiple endotypes^11^. Crucially, it is plausible that only certain endotypes within a given phenotype may respond to a specific treatment^11^.

**Figure 1.**
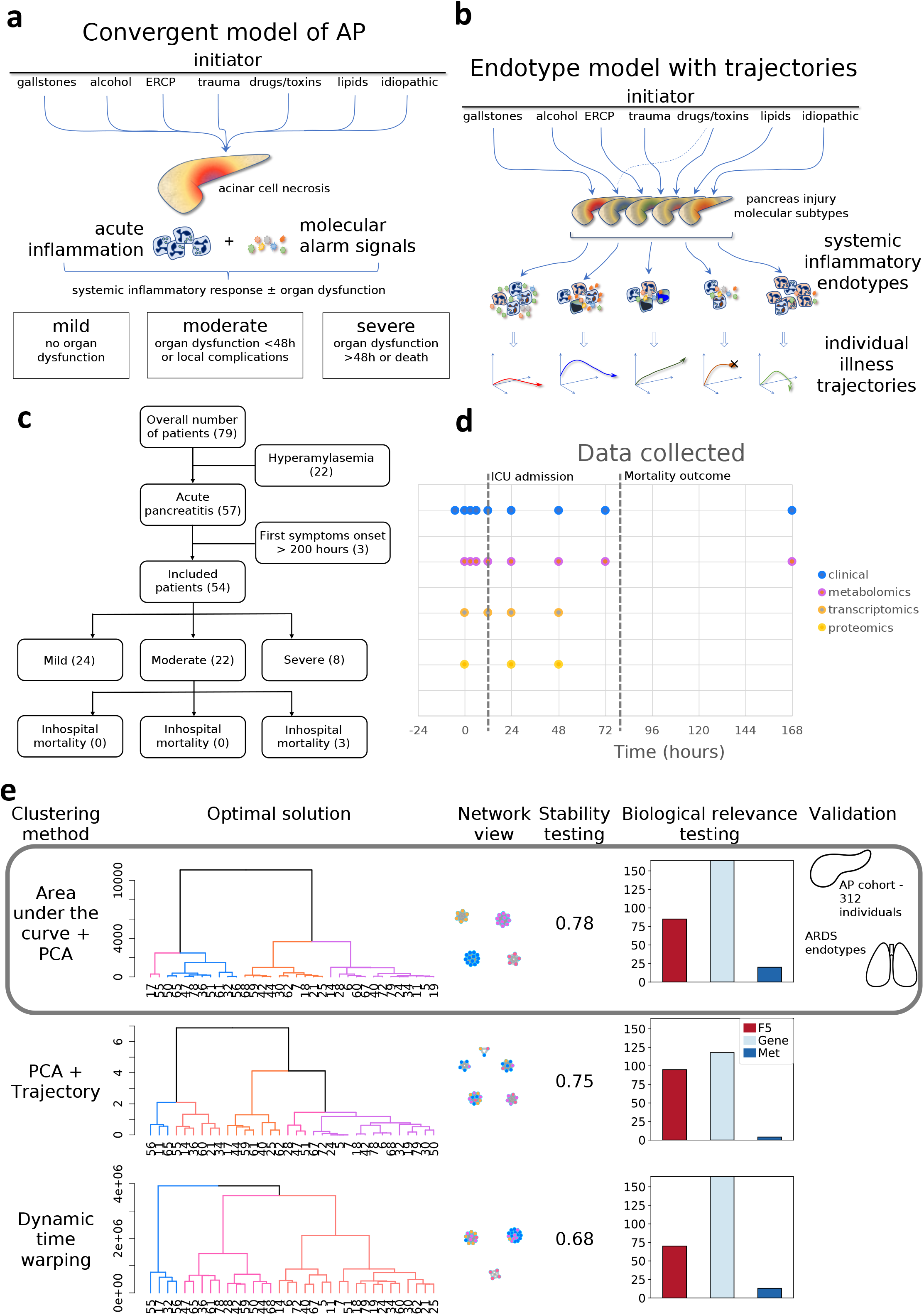
Study concepts, design and discovery of AP endotypes. **A**. Convergent model of acute pancreatitis where diverse etiologies trigger an initial acinar cell damage that results in systemic inflammation that can be mild, moderate or severe. **B**. Endotype model. Proposed acute pancreatitis model suggesting the existence of clinically and therapeutically relevant molecular endotypes. **C**. Study flow chart for patient samples and data from the IMOFAP study included in the analysis showing filtering process and reasons for exclusion. **D**. Sample and data time points. Dashed lines show the median time from admission to hospital to intensive care admission in those who required it (12 hours) and median time from admission to hospital to death for AP fatalities (82 hours)^7^. **E**. Pipeline overview using the 34 pre-selected IMOFAP individuals (individuals with less than 2 time points were not included in the analysis, n=20). Hierarchical trees for each time series based clustering method are presented along with the optimal solution. Colours used to represent the nodes in the network view reflect the 4 groups obtained using the AUC combined with PCA method. Each of the clustering stability measures is reported (average Jaccard index) and a summary of the number of pathways differentially expressed is shown for each category (respectively FANTOM5 results, gene-based results and metabolic compound results). For each one of the three methods based on time series, the best solution, equivalent to the optimal number of clusters (choice based on highest Jaccard index), is presented along with stability and pathway analysis results summary. Results based on a single time point (using Euclidean distances) were not presented here as the clusters obtained using these presented poor structure (mostly a main group with the majority of individuals and a number of single-patient groups).

Until now, evidence for the existence of molecular endotypes in AP remained elusive. To address this, we integrated co-incident peripheral whole blood RNA-sequenced transcriptomic, 10-plex tandem mass tag (TMT) plasma proteomic, and serum metabolomic data, comprehensively annotated with contemporaneous clinical and physiological data, obtained at serial time points from n=54 patients from the IMOFAP cohort (Inflammation, Metabolism and Organ Failure in AP)^12^, of which n=24 had mild AP, n=22 had moderately severe AP, and 8 had severe AP requiring critical care, and the overall number of deaths was n=3 (**Figure 1c and Supplementary Table 1**). Because the majority of patients who develop AP-MODS are admitted to critical care within 48 hours after presentation to hospital^7^, we included serial time points between 0 and 48 hours after presentation (**Figure 1d**). We took an open approach to analysis, without preconceived notions of expected dominance of certain molecular mechanisms, and initially included all possible variables from the data to avoid bias due to previous findings or hypotheses. The total starting data set consisted of the relative expression of a normalised set of 19766 genes from 75bp paired-end Illumina reads RNA-Seq data, integrated with 1383 protein abundances obtained using 10-plex Tandem Mass Tag (TMT), and abundances of 686 identified metabolites, curated according to containerized standards and are made openly available (link to DataStore). After pre-processing, we created a combined data file consisting of 651 metabolites, 371 proteins and 19766 genes that was used as input.

To discover AP endotypes, we applied unsupervised clustering strategies to patient-to-patient distance matrices, as in our previous work^13^. Specifically, we computed single time-point Euclidean distances, area-under-the-curve (AUC) values within a principal component analysis (PCA) space, state-space trajectories in PCA space^14^ and dynamic time warping (DTW)^15^ distances. Once dissimilarity matrices were obtained, hierarchical clustering with the agglomerative Ward’s method^16–18^ was applied to create dendrograms and obtain clusters (**Figure 1e**). The stability of clusters was assessed by bootstrapping, using a 100 iterations with a random hold-out set and quantified using a Jaccard index (JI) of 0.5 as the dissolution point and a JI greater than 0.75 to define stability ^19^. These stability criteria were met by the AUC in PCA space method (JI = 0.78) and state-space trajectories in PCA space (JI = 0.75), demonstrating that clustering generates internally-robust groups (**Figure 1e**). Comparisons with results obtained with the other described methods, for a same number of clusters, showed some similarity (JI = 0.53 when comparing with dynamic time warping results and JI = 0.25 when comparing with trajectory results as illustrated in **Supplementary Figure 1, Supplementary Table 2 and Supplementary Table 3**).

Biological external validity was best for the AUC in PCA space clustering method over alternatives, based on pathway analysis using generalised linear model and the highly-annotated machine-readable KEGG pathway database as the biological gold standard^20^ (extracted from both R libraries GAGE^21^ and MetaboAnalystR^22^) and co-expression clusters derived from the FANTOM5 project^23,24^ as another biological input set. The number of FANTOM5, KEGG genes based and KEGG metabolites based hits for each clustering method is shown in **Figure 1e**. For the AUC+PCA method, the total percentage of variance explained by the first two components used was 51.5%, comprising 40.2% for PC1 and 11.3% for PC2. For this four-group solution, the silhouette score^25^ based on the distance matrix was 0.39 (range 0.23 to 0.57) (**Figure 1e, Supplementary Table 4**).

If the maximum clinical utility is to be obtained from endotype assignation in AP, the endotype should be identifiable as close as possible to the time the affected individual seeks medical help. Using baseline data only, we computed variable importance in projection (VIP) scores using a partial least square discriminant analysis algorithm (PLS-DA) applied to the AUC+PCA 4-cluster grouping, as shown in **Supplementary Figure 2, Figure 2a and Supplementary Figure 3**. Critically, although the PLS-DA model was built using data solely from baseline, the clusterings obtained using only time = 0 data produced inferior results, with Jaccard indexes never exceeding 0.75 (**Supplementary Figure 4**). This data show that a dynamic (time series) dimension is beneficial for training an illness trajectory model.

**Figure 2.**
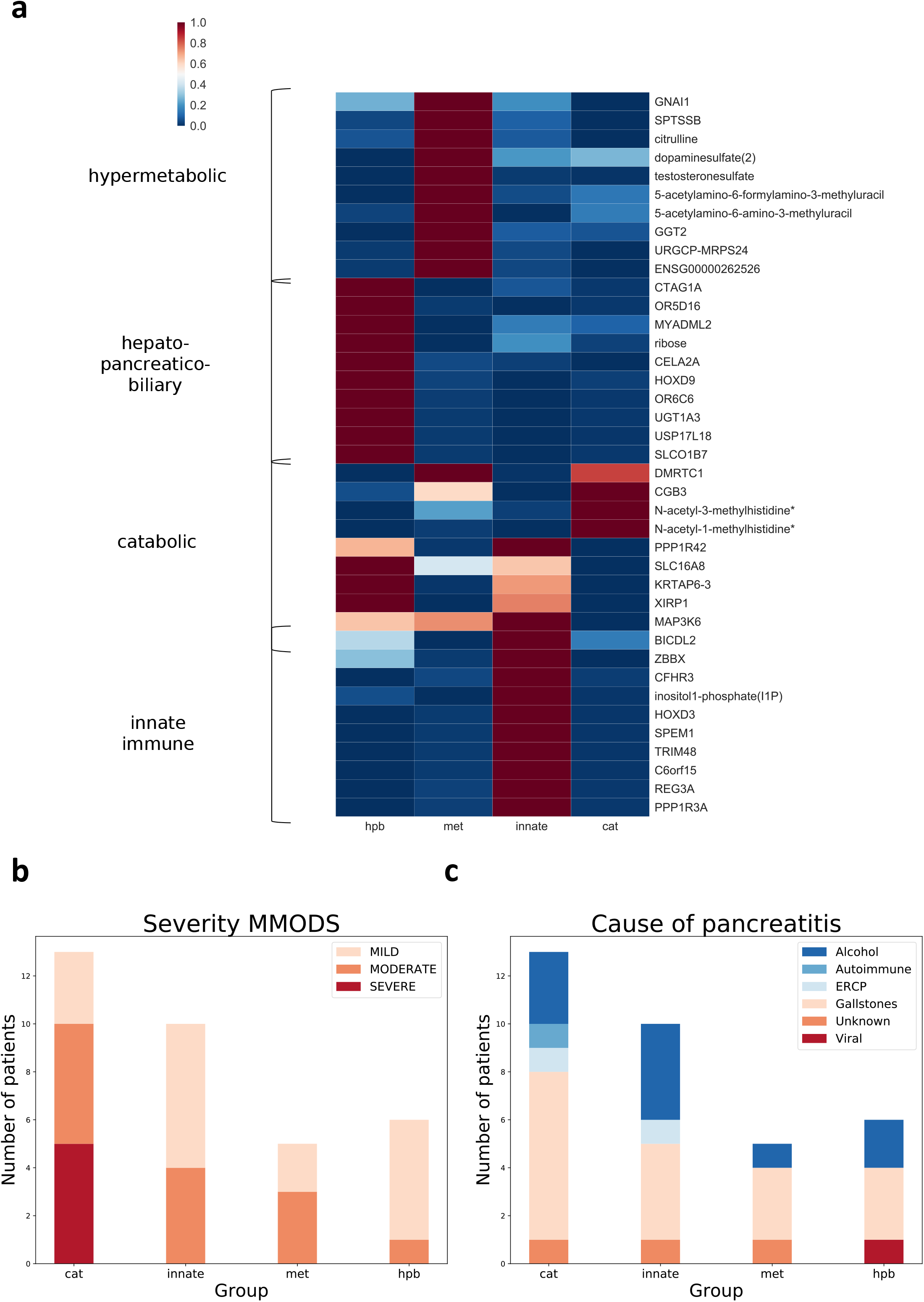
AP identified endotypes. **A**. AP endotypes. The top 10 variables, average values (normalised and scaled) for each identified group are displayed. For visualisation purposes row values were scaled between 0 and 1. Colours are representative of the range of values observed. Values were clustered based on expression patterns considering average values per variable per group. **B**. For comparison purposes, distribution of clinical severity categorised by mMODS score in each endotype. **C**. Distribution of etiology in each endotype. For each endotype, the number of patients is shown.

For each one of the four endotypes we identified the most prominent pathobiological theme and could allocate a name: i) hypermetabolic, ii) hepatopancreaticobiliary (HPB), iii) catabolic and iv) innate immune (**Figure 2a and Supplementary Table 5**). This was done rationally: each identified variable was cross-referenced with the following publicly available online resources GeneCards (Weizmann Institute of Science), HUGO Gene Nomenclature Committee, NCBI EntrezGene, UniProtKB, Ensembl, NCBI PubChem, NCBI PubMed and Google Search, and semantic patterns were identified, discussed and agreed among the authors. Discernable features for the hypermetabolic endotype include GGT2 (g-glutamyl transferase 2) – glutathione homeostasis; GNAI1 (glucosamine 6-phosphate N-acetyltransferase); the caffeine metabolite products of arylamine N-acetyltransferase, 5-acetylamino-6-(+/-formyl)amino-3-methyluracil^26^; dopamine sulphate – a marker of increased gastrointestinal metabolism of endogenous dopamine^27^; citrulline – integral to the tricarboxylic acid cycle; and SPTSSB (serine palmitoyltransferase subunit B) – the rate-limiting enzyme for sphingolipid biosynthesis^28^. For the HPB endotype, the thematic features were CELA2A – pancreatic elastase 2; UDP-glucuronosyltransferase – which is associated with Gilbert-type hyperbilirubinemia^29,30^; and SLCO1B7 – a liver-specific organic anion transporter involved in bile secretion. The catabolic endotype was named for the following prominent features: N-acetyl-3-methylhistidine and N-acetyl-1-methylhistidine – increased after muscle myofibrillar proteolysis and in renal failure^31^; SLC16A8 – a lactate and ketone body transporter; XIRP1 – encoding Xin, a muscle-specific actin binding protein upregulated within 12 hours of injury^32^; and MAP3K6 – a mitogen-activated protein kinase kinase involved in apoptosis signalling^33^. The innate immune endotype defining features were: complement factor H-related protein – a heparin-binding protein involved in complement regulation; inositol-1-phosphate – the basis of inositol signalling; HOXD3 – upregulation of which increases immune cell adherence by upregulating glycoprotein IIb/IIIa^34^; TRIM48 – integral to interferon-*γ* signaling and oxidative stress-responsive cell death via apoptosis signal-regulating kinase 1^35^; PPP1R3A – which has a genetic association with type 2 DM and familial partial lipodystrophy 3^36^; and REG3A – which encodes a bactericidal C-type lectin known commonly as pancreatitis-associated protein that, among multiple actions, alters the gut microbiome and regulates gastrointestinal inflammation^37^ (**Figure 2a**). Our endotype nomenclature was also supported by a pathway analysis. In view of this evidence of distinct functional or pathophysiological mechanisms, we conclude that these groups represent disease endotypes as defined in the Stratified Medicines Framework^10^.

All participants with AP-MODS clustered in the catabolic endotype, despite data on clinical outcome being initially withheld from the model (**Figure 2b**). Although this finding was statistically significant (severe vs. non-severe, Fisher-Freeman Halton test, p=0.038), it is too early to determine whether this would be replicated in a larger cohort. However, the etiology of AP was distributed evenly across endotypes (Fisher-Freeman Halton test, p=0.97) (**Figure 2c**). Gender (Fisher-Freeman Halton test, p = 0.67) or time of onset of symptoms (ANOVA, p = 0.97) were not statistically significantly associated with endotype. Moreover, we generated a clustering for the AUC combined with PCA method using the residuals from a linear model which included gender, age and time of onset. The 4-cluster solution obtained using the corrected data was compared to the chosen partition and showed a high level of similarity (JI = 0.82 and distance matrices correlation, using a Mantel test, showed a correlation of 0.91 with an associated p-value of 0.01). Independence between endotypes and systemic inflammatory response syndrome (SIRS) was tested for and was not rejected (SIRS vs no SIRS, Fisher-Freeman Halton test, p=0.097) (**Supplementary Figure 5**).

To externally validate the generalisability of our findings, we used our results in an independent dataset of AP patients, the KAPVAL cohort (**Figure 3a**). KAPVAL (n = 312 patients, **Supplementary Figure 6, Supplementary Table 6**) has clinical (including laboratory results) and metabolomic data obtained at the time of presentation to hospital with AP, and outcome data regarding death, critical care admission and duration of hospital stay. Reported deaths were not independent from group allocation (Fisher-Freeman Halton test, p < 0.001, **Figure 3b**). Admission to critical care (ICU/HDU vs. ward stay only) was also dependant of group allocation (Fisher-Freeman Halton test, p < 0.001, **Figure 3c**). Length of stay varied between the groups with the reported values of the catabolic endotype globally higher when compared to others (median values were 6.25, 5.13, 4.33 and 4.63 days respectively, for each group with corresponding interquartile ranges 7.25, 10.92, 5.46 and 4.67 days, and Q1-Q3 3.25-10.54, 2.46-13.42, 1.46-6.92 and 3.38-8.00 **Figure 3d**). Average values per time point per group for each one of the identified endotypes are presented in **Supplementary Figure 7** for a subset of variables. For all compounds with a VIP associated value greater or equal to 2 averaged time profiles can be obtained via an online page that can be accessed through the following address: http://baillielab.net/pancreatitis/ (username: pancreas and password: review) along with some clinical and cytokine measures.

**Figure 3.**
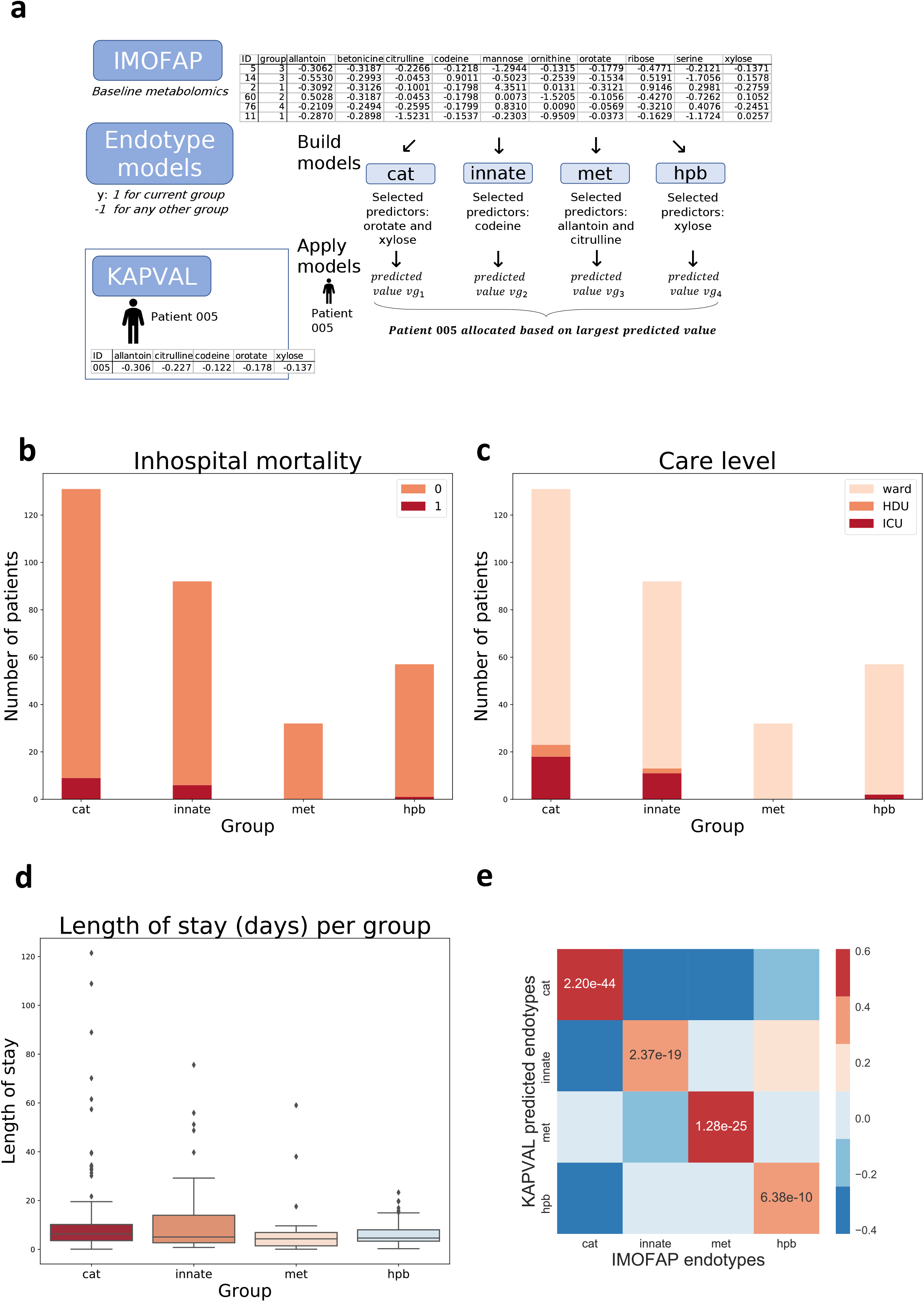
Internal validation of AP endotypes. **A**. Schematics representing the process to assign KAPVAL individuals to endotypes identified in IMOFAP cohort. **B**. Distribution of in-hospital mortality in each endotype for KAPVAL individuals. 1 refers to death and 0 to no death. **C**. Distribution of care level in each endotype for KAPVAL individuals. D. Box plot of length of hospital stay (in days) per identified endotype within KAPVAL cohort. Bars represent 95% confidence intervals. **E**. Correlation matrix representing Spearman’s correlation results for pairwise comparisons between ranked variables from training set (IMOFAP) and testing set (KAPVAL). P-values associated to correlation coefficients computed between matching IMOFAP and KAPVAL predicted groups are shown.

We computed Spearman’s correlation (comparison of ranks, validated using t-distribution) for each pairwise comparison of groups from both cohorts, using metabolites that were not used to predict the group allocations (**Figure 3e**). We obtained significant results when comparing groups from IMOFAP cohort to their corresponding groups in KAPVAL cohort (correlation coefficients ranged from 0.29 to 0.61, corresponding p-values 2.20E-44, 2.37E-19, 1.28E-25 and 6.38E-10 for each endotype, respectively). This external validation demonstrated underlying biological similarity between corresponding endotypes and was unlikely to be observed by chance.

Additionally, we noticed a similarity between our endotype analysis and a disease endotype reported in a related but distinct condition, acute respiratory distress syndrome (ARDS)^38^. Severe AP can cause ARDS, but there are multiple other causes of ARDS including sepsis, trauma, and major surgery^39^. We hypothesised that the hypermetabolic AP endotype reflects the same underlying disease process as the “Type 1” ARDS endotype reported by Calfee *et al*^35^. In order to test this hypothesis we matched measurements from our dataset with the measurements used to define ARDS endotypes. We found 20 variables that matched the 31 variables in the ARDS study (6 physiological, 7 clinical biochemical, and 2 cytokine variables that were not used to produce the clusterings, and 5 genes that were). There was a significant negative correlation (Spearman’s) between the catabolic AP endotype with phenotype 1 in the two ARDS cohorts reported elsewhere (ALVEOLI p = 0.004; ARMA p < 0.001). Interestingly, the hypermetabolic AP endotype correlated positively with the ALVEOLI (p = 0.006) and ARMA cohorts (p < 0.001). (**Figure 4a and Figure 4b**) using AVEOLI ordered variables. This provides compelling additional external validation for our observations and demonstrates that the endotypes or AP that we report here are generalisable to another type of critical illness.

**Figure 4.**
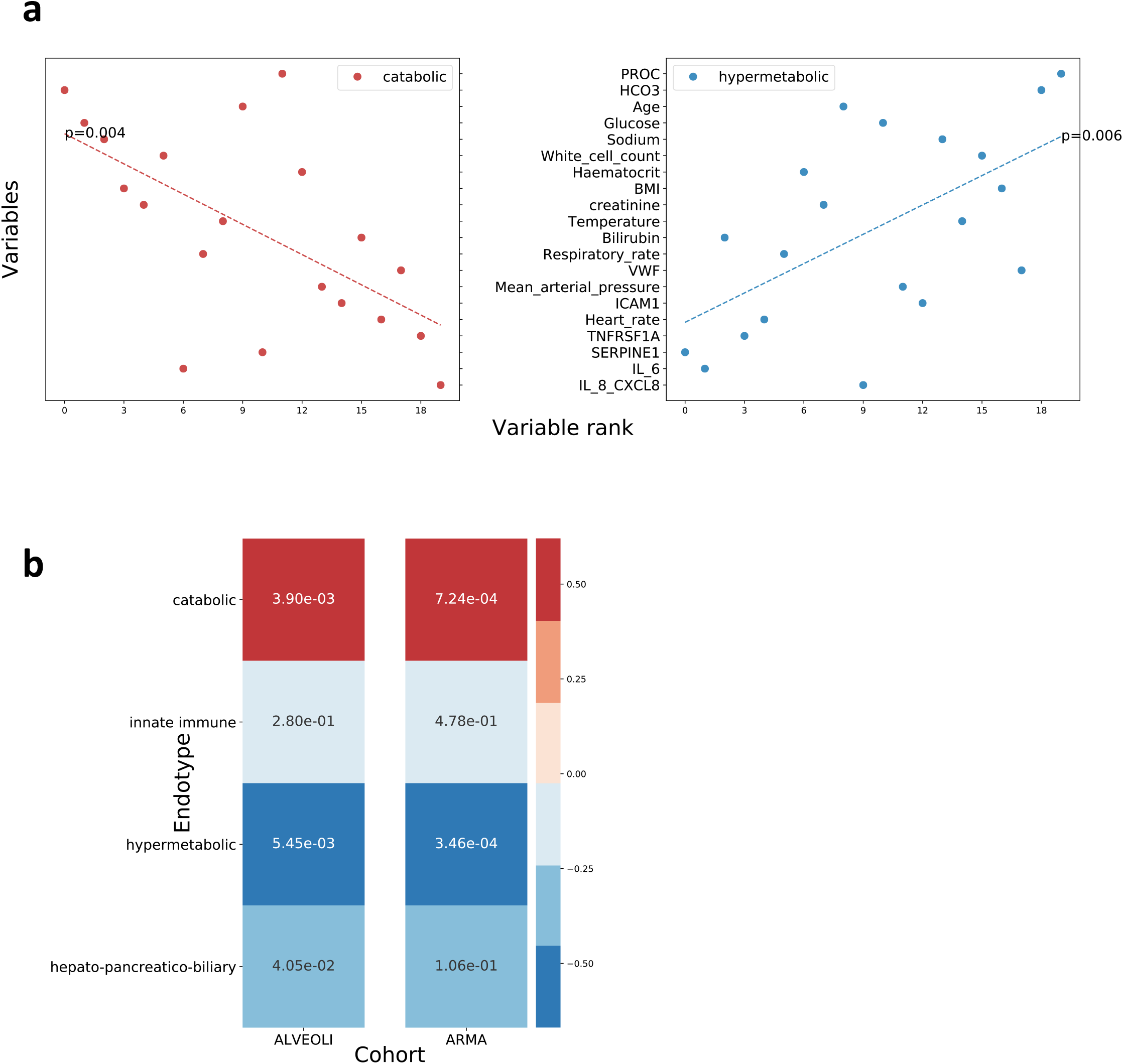
External validation of AP endotypes. **A**. Ordered normalised values represented for catabolic and hypermetabolic endotype. Variables that occur in common with those reported in the ARDS study of Calfee *et al* are presented. Linear trends and ranks of variables based on average value per variable per group were generated and represented using ALVEOLI ordered variables. **B**. Spearman correlation coefficients between identified groups and ARDS cohorts. P-values are reported for all pairwise comparisons.

Finally, we highlighted a significant overlap (Jaccard index 0.88) between our proposed endotypes and groups highlighted using a multi-omics data integration framework, MOFA^40^, when using AUC values. We also noticed a great difference in structure between our endotypes and the ones highlighted using MOFA when applied to baseline KAPVAL data, confirming that, baseline data alone cannot be used directly to highlight the structure of AP endotypes and that previous knowledge of patient trajectories is required to generate the endotype models. Once established the models could be applied to presentation data.

In summary, our data confirm the existence of pathobiological mechanism-based endotypes in AP that could not otherwise be described using current clinical measures of severity or etiology. Using a novel serial evaluation approach, we identify four endotypes that passed stability and biological relevance validation, and that can be characterized by key pathways and biomarker combination stereotypes into hypermetabolic, hepatopancreaticobiliary, catabolic and innate immune endotypes. These AP endotypes are statistically stable and externally validated. Our findings represent a step-change advance in the understanding of acute pancreatitis and lay the foundation for prospective identification of individuals by endotype who may respond differentially to novel therapeutics as these emerge from the translational pipeline.

## Materials and Methods

### Ethics & Regulatory Approvals

The IMOFAP study was approved by the Scotland A Research Ethics Committee (ref: 13/SS/0136 and amendment AM01), NHS Lothian Research & Development Project Number: 2013/0098 and amendment SA1. The University of Edinburgh and NHS Lothian ACCORD were sponsors. NHS Lothian Caldicott Guardian permission was obtained for access to identifiable patient data where needed prior to linked anonymisation. Adults without the capacity to give informed consent were recruited in accordance with the Adults with Incapacity (Scotland) Act 2000, Part 5, through their legal representative, and informed consent was sought when capacity was regained. All participants with capacity provided written informed consent. The study was registered on the UK Clinical Trials Gateway (ref: 16116). The KAPVAL cohort was approved by the East of Scotland Research Ethics Committee (ref: 15/ES/0094) and NHS Lothian Research & Development Project Number: 2015/0447/SR594, under the aegis of the Lothian NRS Human Annotated Bioresource. The University of Edinburgh and NHS Lothian ACCORD are sponsor. NHS Lothian Caldicott Guardian permission was obtained for access to identifiable patient data where needed prior to linked anonymisation.

### Clinical sample and routine laboratory data acquisition

We used samples and clinical data from the IMOFAP cohort, which has been described in full previously^12^. In brief, consecutive emergency attendees to hospital with sudden onset abdominal pain with nausea and/or vomiting, and a serum amylase value greater than 100 iU/L were identified using an automated laboratory alert system with clinical verification, at all times of day or night for a period of three months. Confirmation of the diagnosis of AP according to the revised Atlanta criteria^41^ was made after recruitment in 57 of 79 recruited patients. For the integrated multiomic analysis presented here, we applied a further exclusion criterion to 3 patients who had a prolonged interval (greater than 200 hours) after symptom onset to first study sample, because we decided that those patients would have been likely to enter this study at a late point on their individual disease trajectory and thus introduce bias (**Figure 1c**). The median time interval between symptom onset and recruitment in the 54 included patients was 21.3 hours (interquartile range 40.8 hours, Q1-Q3 13.5-54.4). Timed samples of peripheral venous blood were taken at recruitment and 3, 6, 12, 24, 48, 72 hours after that, and again at seven days. Samples were aliquoted into specific tubes containing appropriate preservation solution for subsequent DNA and RNA extraction, or serum and plasma extracts were prepared after centrifugation and snap frozen (**Figure 1d**). Although every effort was made to capture all time points, this was not always possible, and on occasion (for example, at the request of the patient to omit a short interval repeat venepuncture), time points were omitted. From this biobank, samples were selected for analysis based on the completeness of the multiomic set for a single individual. When analysing single time point data, and more precisely time point 0, we selected patients based on whether or not they had a complete multiomic set at time point 0, leaving 40 of the pre-selected 54 patients (consisting of 22 mild, 14 moderate and 4 severe AP cases). Differently, when comparing time-series we required at least two complete time points, selecting 34 patients (including 16 mild, 13 moderate and 5 severe AP cases).

We used samples and data from n = 312 patients from the KAPVAL cohort for validation. The KAPVAL cohort is a fully-linked anonymised surplus biosample cohort which is made up of samples and data from all patients presenting to the Royal Infirmary of Edinburgh with a serum amylase level > 300 iU/L (3-fold above the upper limit of the reference range for our laboratory). An aliquot of gel-clot activator serum is retained and stored at −80 °C for all patient samples that have an elevated serum amylase. Using a linked anonymisation code, to which the investigators are blinded, the diagnosis of acute pancreatitis was confirmed by trained members of a specialist data collection team, using clinical and laboratory data obtained from the individuals electronic health record. The diagnosis of AP was made according to the revised Atlanta criteria^41^. Again, this data is not shared with the research team until it has been fully anonymised and all personal identifying information removed. A unique study identifier is used to subsequently link the serum sample with the clinical annotation prior to providing these to the investigator team. The variables used in the analysis include age, gender, date and time of admission and discharge, first 3 diagnosis codes, standard clinical biochemistry and clinical haematology tests, level of critical care, duration of critical care and mortality. Serum samples were used as described.

### Transcriptomic data acquisition and pre-processing

2.5 ml of peripheral venous blood was collected into the PAXgene Blood RNA Tube (BD Biosciences) following the manufacturer’s instructions and stored at −80 °C until used. Total RNA was extracted and purified using the PAXgene Blood miRNA Kit (QIAGEN). The RNA integrity of total RNA samples was assessed using the Agilent 2100 Bioanalyzer. The mRNA in a total RNA sample was converted into a library of template molecules of known strand origin using the reagents provided in an Illumina^®^ TruSeq^®^ Stranded mRNA library prep workflow. The subsequent sequence data was obtained using Illumina HiSeq 4000 75PE system. RNA-Seq data consisted of 75bp paired-end Illumina reads stored as FASTQ files. One batch was carried out using polyA selection and the other using rRNA depletion. Samples were filtered based on QC results (FASTQC). Read alignment was performed against the genome assembly hg38 using STAR^42^. Counts were generated as a proxy for gene expression by assigning previously aligned reads to exons using the tool featureCounts^43^. hg38 genome was used as the reference genome. The difference in RNA sequencing (library preparation) was accounted for using a protein-coding only filter, a batch removal algorithm (using the ARSyNseq function from NOISeq R library^44^) and a normalisation step (FPKM). Finally, the normalised counts were transformed into Z-scores. This allowed a comparison across samples. PCA plots of the counts before and after batch effect removal are available in **Supplementary Figure 8**.

### Proteomics data acquisition and pre-processing

Serum was obtained from peripheral venous blood by centrifugation and stored at −80 °C until used. Sera were subjected to depletion of abundant serum proteins using Proteome Purify 12 Human Serum Protein Immunodepletion Resin (R&D Systems). Denaturing was followed by alkylation with N-ethylmaleimide and acetone precipitation. Digestion used lysyl endopeptidase (LysC) and trypsin before labelling with 10plex TMT reagents (Thermo Fisher Scientific). TMT-labelled peptides were fractionated into 4 fractions each by High-pH Reverse Phase chromatography then each fraction analysed by RPLC-MS/MS/MS (70 min linear gradient) on a Fusion Tribrid Orbitrap operating in Data Dependent Acquisition mode (MultiNotch Simultaneous Precursor Selection method; MS1: profile mode, Orbitrap resolution 120k, 375-1550 m/z, AGC target 200,000, 50 ms max. injection time, RF lens 60%; MS2: centroid mode, IonTrap, 12 dependent scans, 1.2 Th isolation window, charge states 2-7, 60 s dynamic exclusion, CID fragmentation (35%, activation Q 0.25), AGC target 10,000, 70 ms max. injection time; MS3: profile mode, 5 precursors, 2 Th isolation window, Orbitrap resolution 50k, 100-500 m/z, AGC target 50,000, 105 ms max. injection time, HCD fragmentation (60%)). Control samples were used as internal cross-channel controls in different TMT samples and in different TMT channels to avoid any specific bias. Raw files were searched with MaxQuant (version 1.5.7.4) against a human proteome obtained from UniProt, with the match-between-runs option selected to allow for transfer of peptide identifications between files. After raw data acquisition and initial processing to generate intensity-based values, any protein species with 90% or more missing values was discarded. Scaling and linear imputation were applied using the minimum value for each compound, as any missing value would suggest a value below the detection limit. As samples from similar batches grouped together when performing the clustering step, we corrected for it using ComBat to remove the irrelevant variation between samples due to the different runs carried out. Measurements were then transformed into Z-scores.

### Metabolomics data acquisition and pre-processing

Serum was obtained from peripheral venous blood by centrifugation and stored at −80 °C until used. Aliquots of sera were shipped on dry ice to Metabolon Inc., 617 Davis Drive, Suite 400, Durham, NC 27713 USA. Serum samples underwent automated protein depletion using methanol (MicroLab STAR^®^ system) followed by four fraction analysis by UPLC-MS/MS with positive ion mode electrospray ionization, UPLC-MS/MS with negative ion mode electrospray ionization, LC polar platform and, GC-MS. QA/QC steps included: a pooled matrix sample as a technical replicate throughout, extracted water samples as process blanks, and a bespoke cocktail of QC standards spiked into every sample for instrument performance monitoring and chromatographic alignment. Instrument variability was determined by calculating the median relative standard deviation (RSD) for the standards. Overall process variability was determined by calculating the median RSD for all endogenous metabolites (i.e. non-instrument standards) present in 100% of the pooled matrix samples. Ultrahigh Performance Liquid Chromatography-Tandem Mass Spectroscopy (UPLC-MS/MS): The LC/MS portion of the platform used a Waters ACQUITY ultra-performance liquid chromatography (UPLC) and a Thermo Scientific Q-Exactive high resolution/accurate mass spectrometer interfaced with a heated electrospray ionization (HESI-II) source and Orbitrap mass analyzer operated at 35,000 mass resolution. Gas Chromatography-Mass Spectroscopy (GC-MS): The samples for GC-MS was derivatized under dried nitrogen using bistrimethyl-silyltrifluoroacetamide and separated on a 5% diphenyl / 95% dimethyl polysiloxane fused silica column (20 m x 0.18 mm ID; 0.18 um film thickness) with helium as carrier gas and a temperature ramp from 60° to 340°C in a 17.5 min period and analyzed on a Thermo-Finnigan Trace DSQ fast-scanning single-quadrupole mass spectrometer using electron impact ionization (EI) and operated at unit mass resolving power. Data Extraction and Compound Identification: Raw data were extracted, peak-identified and QC processed using Metabolon’s hardware and software and compounds were identified by comparison to library entries of purified standards or recurrent unknown entities. Metabolite Quantification and Data Normalization: Peaks were quantified using area-under-the-curve. Where runs spanned multiple days, a data normalization step was performed to correct variation resulting from instrument inter-day tuning differences. Metabolomics data consisted of an abundance list (raw ion counts). Data were pre-processed using a similar pipeline as the one used for the proteomics data but did not require a batch effect removal step.

### Data Analysis

Analyses were carried out using Python (version 3.5), R (version 3.3.2) and SPSS (IBM, v24). Libraries used include dtwclust in R and numpy, pandas, rpy2, scipy, sklearn in Python. All methods used pre-processed Z-scores as initial input.

#### Single time point Euclidean distances

The first considered strategy consisted of computing Euclidean between all pairs of patients at selected time points, using pre-processed data. This was performed for time point 0, 24 and 48 hours individually and obtained measures were used as a way to quantify dissimilarity between individuals.

#### Area Under the Curve and PCA

For each variable and for all individuals we computed area under the curve (AUC) values for the corresponding time series using the trapezoidal rule and based on pre-processed values as described for each individual datatype. In doing so, each time series was summarised as a single value representing the cumulative magnitude of response over time. Consequently, values obtained for each variable could be treated as independent from others. We normalized the values based on the time difference between the first and last included time points. Using this newly created dataset we projected the values on a principal component analysis plot where, selecting the first two components, we computed Euclidean distances between the different individuals. The Euclidean distances were weighted according to the proportion of variance explained by each principal component so that the distance between two individuals on PC1 axis would weigh more in the final distance compared to the distance on PC2. PCA is a method of choice when encountering high dimensionality data, as much for data visualization as for data analysis, and is hypothesis free^45^. Indeed, PCA is only sensitive to the correlation structure in the data and does not make specific assumptions related to the stratification of the input data.

#### Trajectory through PCA space

As described in another study^14^, trajectories of patients through selected components can be helpful when clustering patients. Here we projected all time points of each individual onto a two-dimensional PCA space and looked at their evolution through this newly defined space. To characterize the trajectory of an individual through this space we considered the direction taken between each pair of time points for these particular patients. We coded this direction with a value of 1, 2, 3 or 4 depending on the direction taken when dividing the space into four quadrants. This was repeated for all patients and eventually a vector of directions was obtained for each patient. The Hamming distance was used to compare the vectors. When taking two vectors of the same length, the Hamming distance is computed by counting, in an element-wise fashion, the number of different elements. The values were then used as dissimilarity measures between patients. This allowed to combine the advantages of PCA and trajectory analysis.

#### Dynamic Time Warping

Computation of distances between patients using dynamic time warping was carried out using the dtwclust package in R. The algorithm then considers each pair of samples. More specifically, for each variable, a matrix is generated and reports the difference in magnitude between all possible pairs of time points. One matrix of dimensions (time series 1 length*time series 2 length) is obtained for each variable. A summary matrix is generated for each patient by summing the individual matrices associated to each variable, element-by-element. The warping consists of finding a trajectory in that matrix that will minimize the distance between the two series. The process will start from the matrix element corresponding to the first points of the two series being compared (in other terms the first element of the summary matrix) and will end when reaching the last series points (corresponding to the last element of the matrix). As this is performed on a summarized matrix, the selected warping represents a consensus alignment minimizing the summed differences in magnitude between the two multivariate time-series. From this warping a final distance is calculated and can be used as a dissimilarity value. We performed linear imputation on time series when time points were not equally spaced or missing. It was preferred as it allowed a fairer comparison when dealing with time series of different lengths/sampling pattern.

#### Clustering strategy

Using the dissimilarity matrices previously obtained we clustered them using hierarchical clustering and Ward’s method. Ward’s method is an agglomerative method and works by minimizing distances within each group. Although it is usually used for Euclidean related distances (which is not the case for all the presented methods) it has been used successfully for other types of distance^16–18^ and produced the best results here when compared with others (in terms of validation results).

The number of clusters was chosen according to the stability of each solution. For all number of clusters ranging from two to twenty we assessed this using bootstrapping combined with the Jaccard index. For a chosen number of clusters *k* we replaced individuals from the initial cohort to create a new input dataset. It was then used to re-perform the clustering 100 times. Each one of the new results obtained was compared to the initial solution given *k* clusters. Each cluster from the *k* generated clusters was compared to the most similar cluster of the initial solution. The Jaccard index computed the overlap between the two and this was averaged for all matched groups that were part of a solution. After 100 repetitions we obtained a value for each *k* number of clusters by taking the average of the averaged Jaccard indexes and the solution with the highest value was chosen as the best one. Twenty was chosen as the maximum number as any greater value would have resulted in many clusters composed of one or very few individuals. This was not desirable as very little information could have been drawn from it. Additionally, we chose not to select for further analyses any solution with one group or more presenting less than three individuals.

#### Validation strategy

As bootstrapping with cluster comparisons produced a measure of stability, we used it to filter solutions based on a Jaccard index threshold of 0.75, meaning that when the initial dataset was changed we would still get a sufficiently similar structure. Pathway analyses were carried out on pre-selected clusters and allowed to assess biological plausibility. To perform the analysis we filtered our pre-processed data to select only variable values reported at time point 0. Compound identifiers were converted when necessary using R package biomaRt. Data used consisted of KEGG pathways extracted from R package GAGE when analysing gene and protein data and from MetaboAnalystR when analysing metabolites. As four groups were identified, the aim was to assess whether or not a subset of compounds (corresponding to a pathway) was associated with the group label. For each pathway this was tested using generalised linear models. A model was fitted to the data for each compound of a corresponding pathway with the group label being the fixed effect and the response variable being the values associated to this gene. To assess the effect of the group on the values, we performed a likelihood test to compare the newly created model to a null model, returning a single p-value. Using Stouffer’s method to combine p-values, we computed a single p-value per pathway. For elements of a pathway, individual p-values were given a weight corresponding to the inverse of the total number of pathways a gene was involved in. This, this prevented overlapping pathways from biasing our results. R function anova with test=“LRT” was used to do the tests. As the gene sets tested for enrichment were the same for each one of the three methods tested to cluster individuals, p-values were used to quantify the biological relevance of a clustering. Each pathway with an associated value under the threshold of 0.05 was counted as differentially expressed and counts obtained for each method were compared to determine the most biologically relevant result. FANTOM5 data clusters were used to compare cell type gene signatures with our groups and thus allowing the discovery of closely involved cell types. The same strategy was applied to determine if a pathway was differentially expressed or not, using the same 5% cut-off.

Results were used as a way to quantify the biological pertinence, have an overall look at the results and select clustering solutions.

#### Enrichment analysis

Partial Least Square Discriminant Analysis (PLS-DA), a classification algorithm, was used to highlight potential biomarkers and processes associated with single groups. The strategy was to create *k* models for *k* groups obtained, each model aiming at the classification of samples to the group of interest or others (regardless of which other group). Given group labels it will project the data onto a new space, given a number of components selected by the user, and then rotate the components to maximize the separation between samples of different groups. Finally, weights can be extracted and a correlation with each variable computed. This is called Variable Importance in Projection and the higher the value the more the variable will have contributed to the classifier model. We filtered variables deemed significant for the classification task using a VIP threshold value of 1 for each one of the models obtaining respectively lists of size 10216, 6584, 9112 and 7037 for groups 1 to 4 (ref. ^46^). The lists were then used to perform pathway enrichment analysis.

To analyse the list produced for each group we first generated a Reactome database using files freely available from the Reactome website (https://reactome.org/download-data, lowest level pathway files). As our variables were of different type (transcriptomics, proteomics and metabolomics) we generated a merged pathway list using different Reactome files. Pathways were then selected for analysis if they had 10 or more compounds and no more than 500 as they were deemed neither robust nor informative. We then used Fisher’s exact test to obtain a p-value per pathway based on the number of matches present in our list and the total number of compounds considered initially to be included in the list of interest. For each list p-values obtained from this test were then corrected using an FDR based strategy and represented applying a threshold of 0.001 (**Supplementary Figure 9**). Following the same strategy, time points 24 and 48 were also analysed and results added alongside the ones obtained for time point 0.

#### Data visualisation

An interactive webpage (available through URL) was created in order to visualise average AUC values and average z-score values over time for the identified endotypes. A subset of variables was selected using VIP scores from the PLS-DA models and a threshold of 2. In order to be able to visualise some of the clinical and cytokines measurements over time we filtered variables based on ANOVA results and a threshold of 0.05.

#### Validation in an independent dataset (KAPVAL)

To validate the groups previously highlighted we used a separate second cohort, the KAPVAL (Kynurenine pathway in AP, VALidation) cohort consisting of 312 individuals with confirmed AP as defined by the revised Atlanta criteria^41^. For these patients, clinical annotations as well as metabolomics data was available for a single time point corresponding to hospital admission. Prior to normalisation, only variables appearing in both datasets were kept. The data was then normalised in the same way as the IMOFAP metabolomics data, quantile normalisation was performed before transforming the values to Z-scores. To see if the same structure could be observed in both cohorts we classified KAPVAL samples using PLS-DA models (with 3 components) created for each one of the IMOFAP group but using solely metabolites that were available for the KAPVAL cohort as well. Admission IMOFAP data points were used as input to generate the models. VIP scores were computed and a threshold of 25 variables was set as a maximum of features to be included in each model. An optimal number of variables between 3 and 25 was selected based on R^2^ values. To classify individuals from the KAPVAL cohort we applied each one of the models to our individual and allocated them to the group from which they were the closest to (**Supplementary Figure 10**). To compare the biology between the groups of the two cohorts we computed average values per variable per group and compared them between the cohorts as inspired by Sweeney and al.^47^. We only compared metabolites that were not included in the PLS-DA models used to classify KAPVAL individuals (355 metabolites). To perform the comparison between average values we computed Spearman’s correlation (comparison of ranks, p-values computed using a t-distribution) for each pairwise comparison of groups from both cohorts (**Figure 3e**). We obtained significant results when comparing groups from IMOFAP cohort to their corresponding groups in KAPVAL cohort (correlation coefficients ranged from 0.29 to 0. 61, corresponding p-values 2.20E-44, 2.37E-19, 1.28E-25 and 6.38E-10 for each group label respectively). This allowed use to say that biology between corresponding groups was indeed similar and unlikely to be observed by chance.

#### Validation in an external dataset

To compare our endotypes to external data we used ARDS endotypes described in another study^35^. From ARDS endotypes we extracted ranking of the variables used and compared it to rankings in each one of our endotypes. As not all ARDS reported variables were available in our dataset, we matched 20 variables that we used to extract and compare rankings. For each one of the two ARDS cohorts, we compared the lists of ranked variables to each one of our endotypes using Spearman’s correlation coefficients. All coefficients were computed using Python scipy module. Significant results were obtained when comparing our catabolic AP endotype with phenotype 1 from both ALVEOLI and ARMA ARDS cohorts (p = 0.004 and p < 0.001). The correlation between our hypermetabolic AP phenotype and ALVEOLI and ARMA cohorts was positive and significant as well (p = 0.006 and p < 0.001 respectively).

#### Comparison with results from an independent tool (MOFAtools)

MOFA^10^ allows to highlight, in a multi-omics dataset, variables explaining variation within a dataset, using factor analysis. Clustering analysis is available as part of the tools suite. We chose to run the analysis using AUC pre-computed values that were previously used to compute the distances within a PCA space and to highlight our endotype groups. Default parameters were used with the additional filter used to drop model factors explaining less than 1% of the variance in all omics. Once the model was built we chose to cluster the individuals using two latent factors and choosing a 4-cluster solution. When comparing our clusters to the ones obtained using MOFA we obtained an overlap of 0.88 as illustrated in **Supplementary Figure 11** (computed using an averaged Jaccard index) and confirming the validity of our clusters.

Using our validation cohort (KAPVAL) and the available metabolomics data, pre-processed as described previously, we tested if the groups we obtained using PLS-DA models trained on IMOFAP data would highlight a similar structure in the dataset. As presented in **Supplementary Figure 12**, the overlap was quite low (Jaccard index value 0.37). We hypothesised that time-series data was required in order to structure the dataset and that some knowledge of the dynamics was required to classify patients using solely baseline data.

## Funding

L.N. holds a Doctoral Training Program PhD studentship from the Medical Research Council. J. K.B is funded by a Wellcome Trust Intermediate Clinical Fellowship (103258/Z/13/Z) and a Wellcome-Beit Prize (103258/Z/13/A), BBSRC Institute Strategic Programme Grant to the Roslin Institute, and the UK Intensive Care Society. D.J.M. is a Senior Clinical Fellow of the Medical Research Council (MR/P008887/1). The IMOFAP study and metabolomics are part of, and funded by, the University of Edinburgh/GSK Discovery Partnership with Academia collaboration. The study was also co-funded by the Wellcome Trust through the Institutional Strategic Support Fund 3 (ISSF3).

## Conflict of interest

No author has any actual or potential conflict of interest with the publication of this paper.

## Supplementary Figures

**Supplementary Figure 1.**
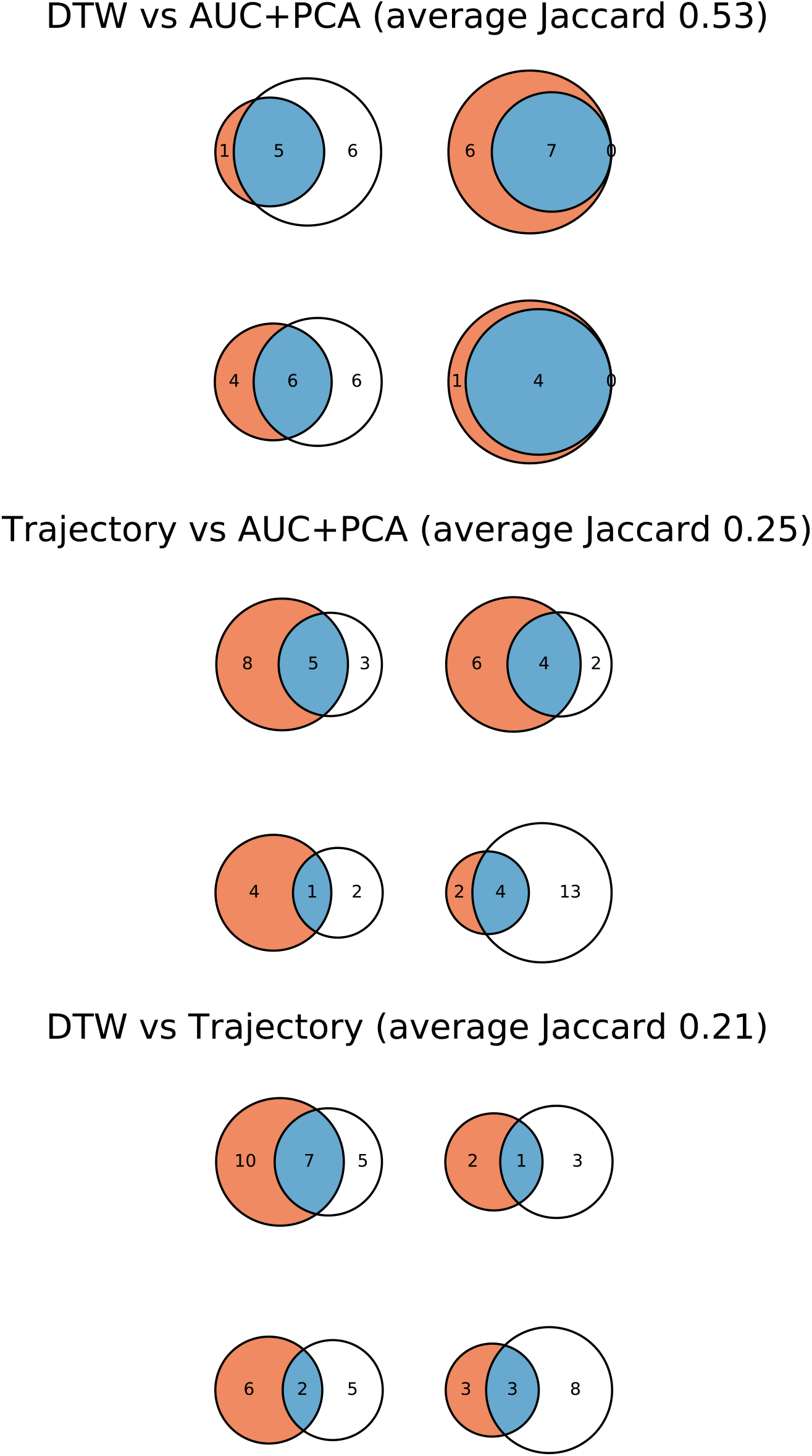
Overlap between the different clustering solutions. For the 4-group selected clustering, comparison with other four-group solutions obtained using the two other methods. Average Jaccard index values are reported for each comparison. Group matching selected based on best overall Jaccard indexes.

**Supplementary Figure 2.**
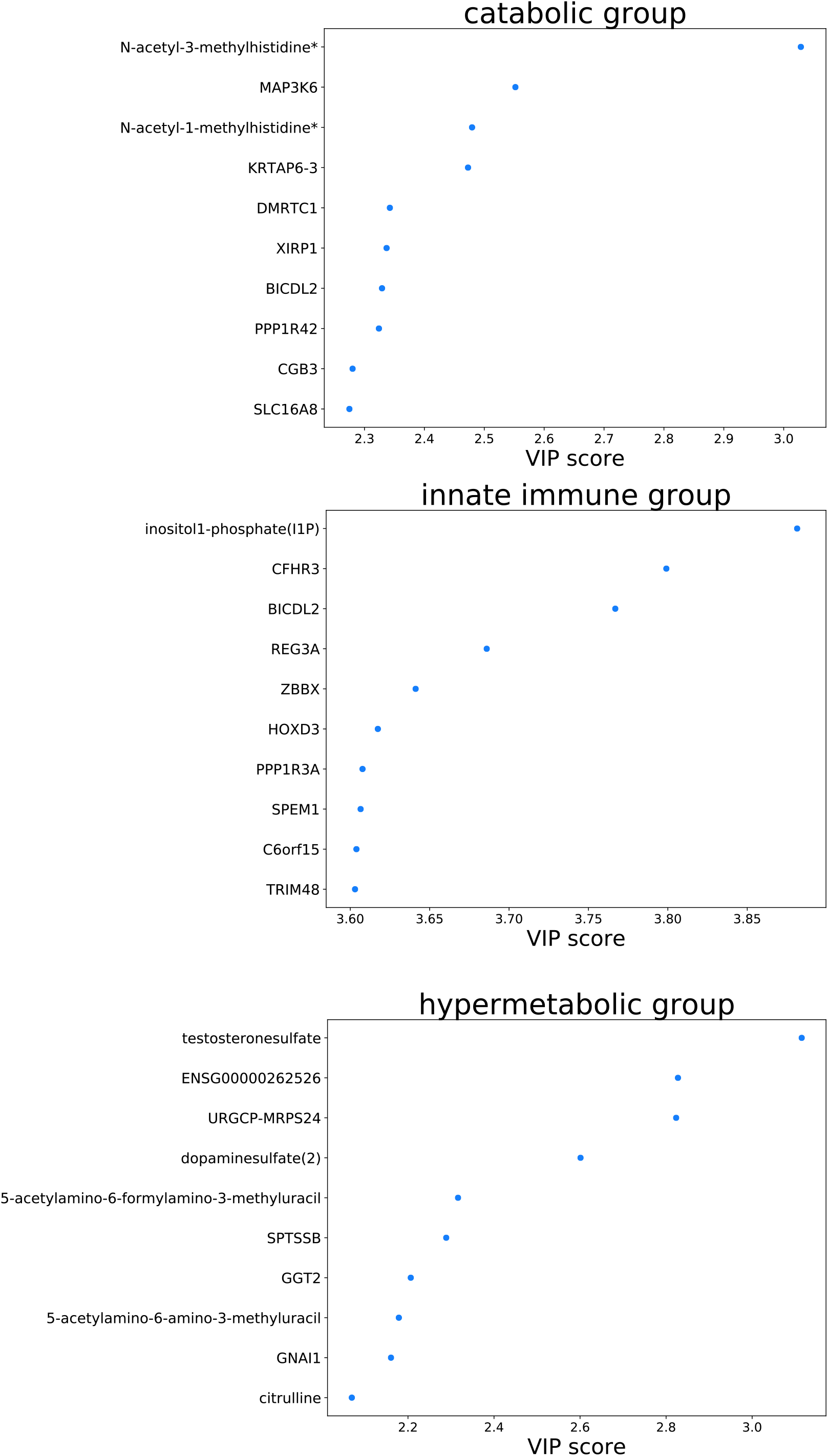
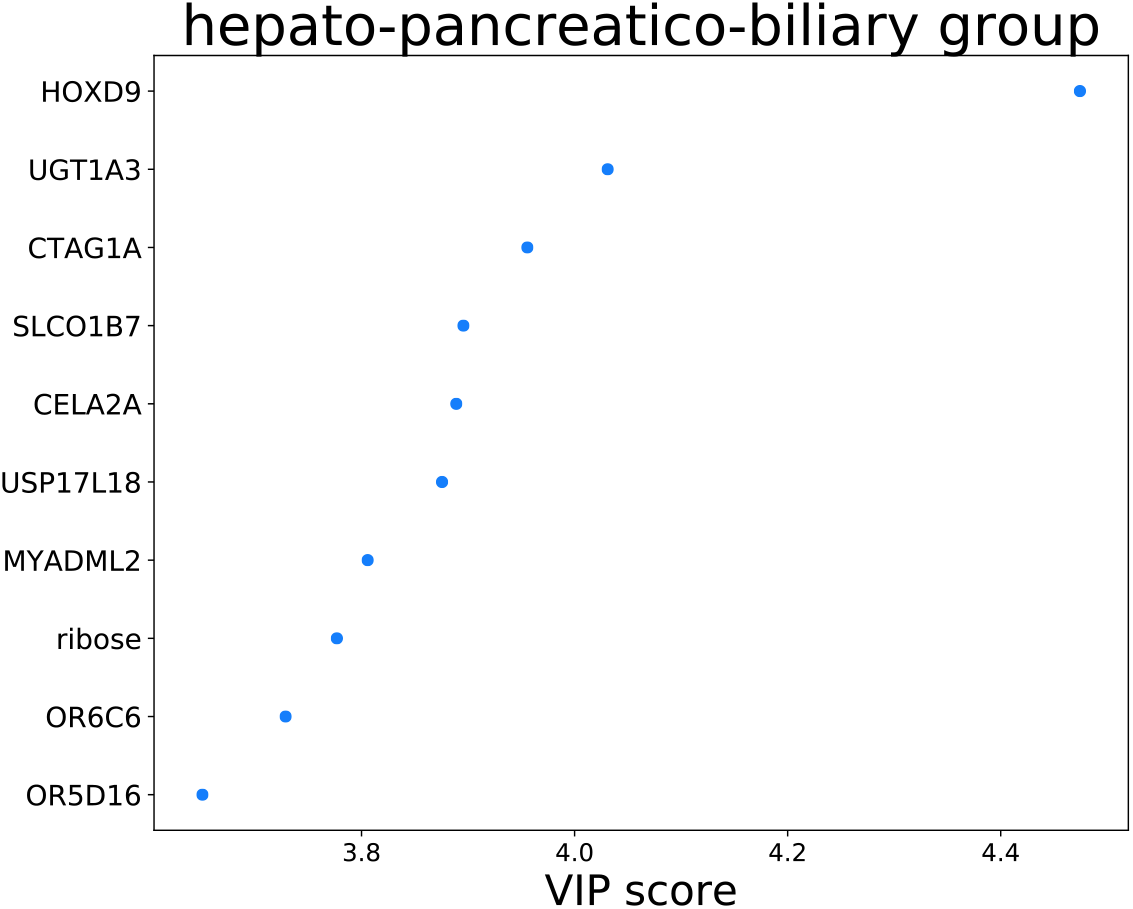
PLS-DA top variables for each identified endotype. The top 10 variables from the PLS-DA model based on VIP values.

**Supplementary Figure 3.**
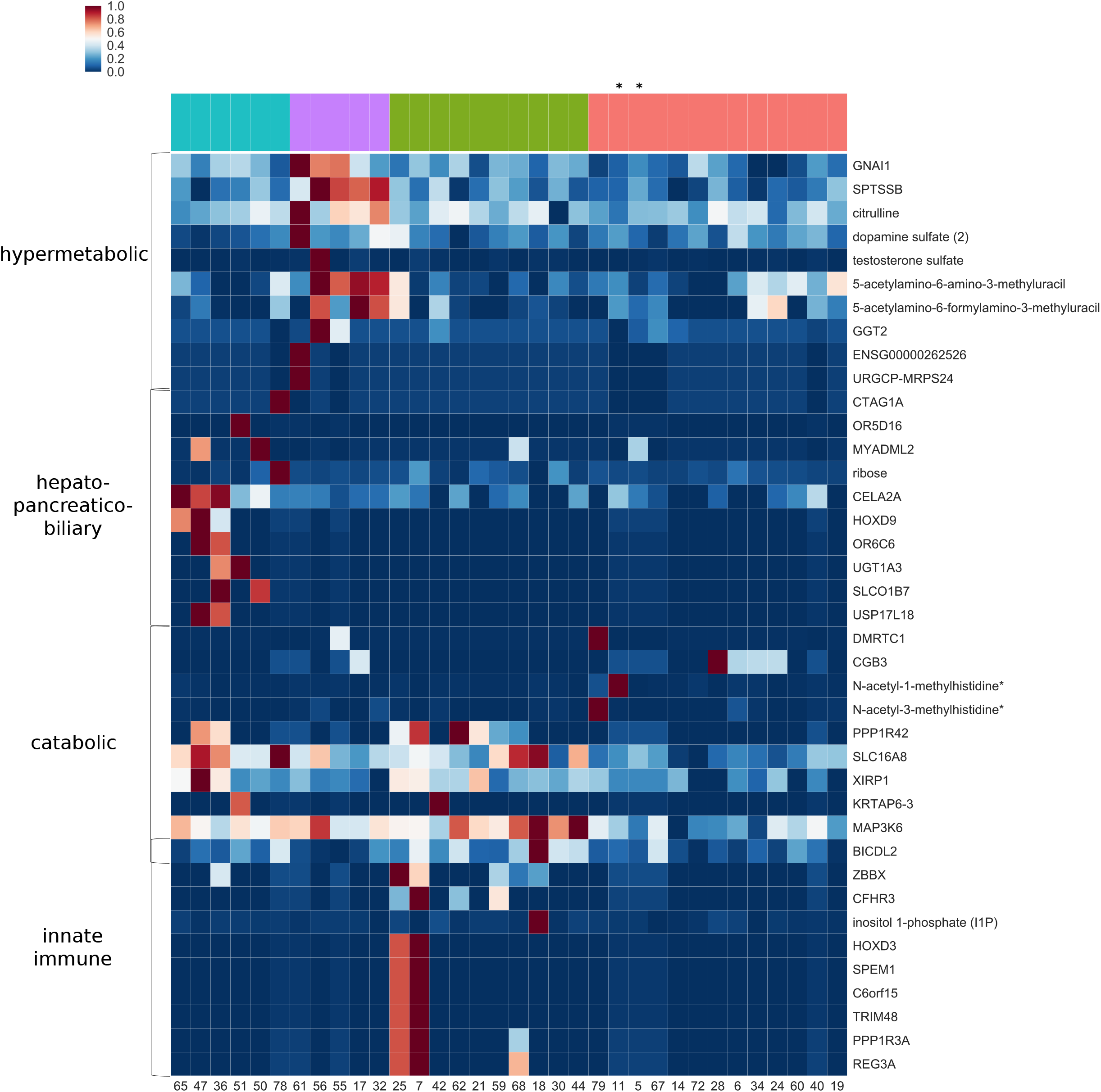
AP endotypes overview for VIP-selected variables. For the top 10 variables, normalised and scaled average values for each identified group are displayed. For visualisation purposes row values were scaled between 0 and 1. Colours are representative of the range of values observed. Values were clustered based on expression patterns considering average values per variable per group. A star indicates a patient who did not survive.

**Supplementary Figure 4.**
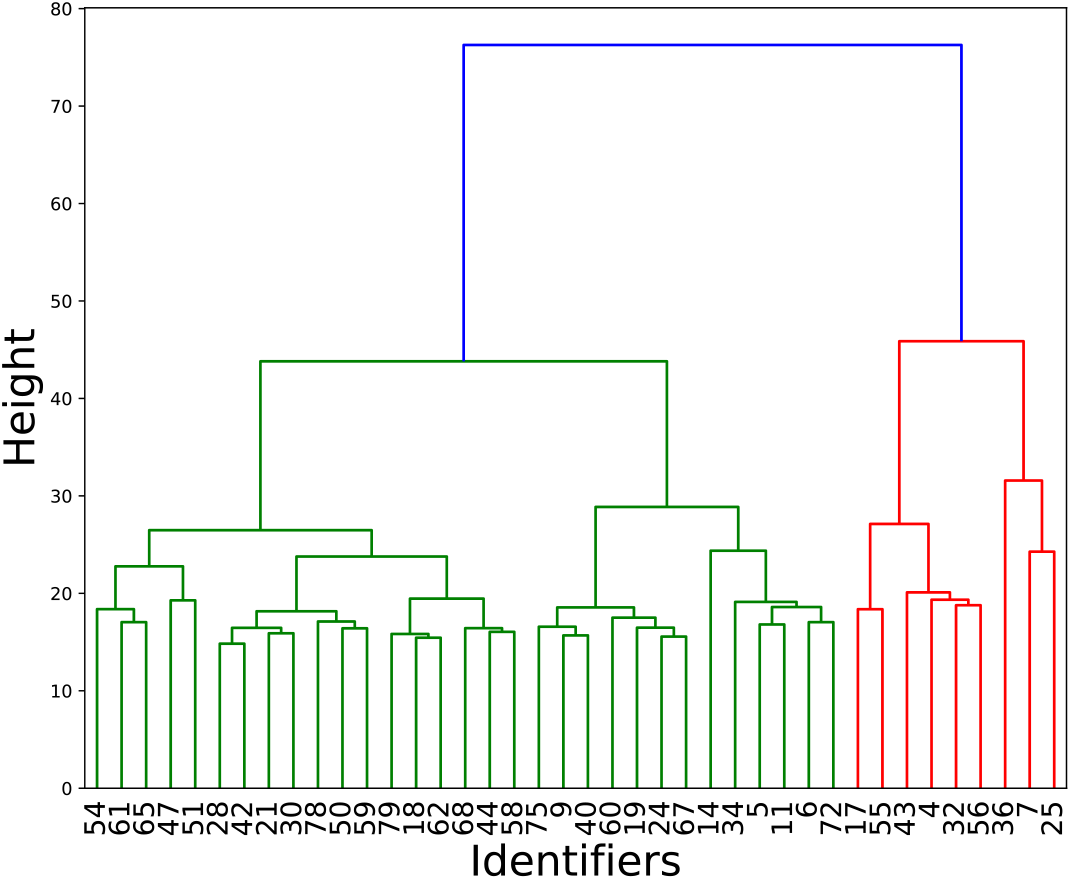
Hierarchical clustering result for time point 0 data. Using only time 0 data and choosing an arbitrary number of clusters, dendrogram based on Euclidean distances and Ward’s algorithm.

**Supplementary Figure 5.**
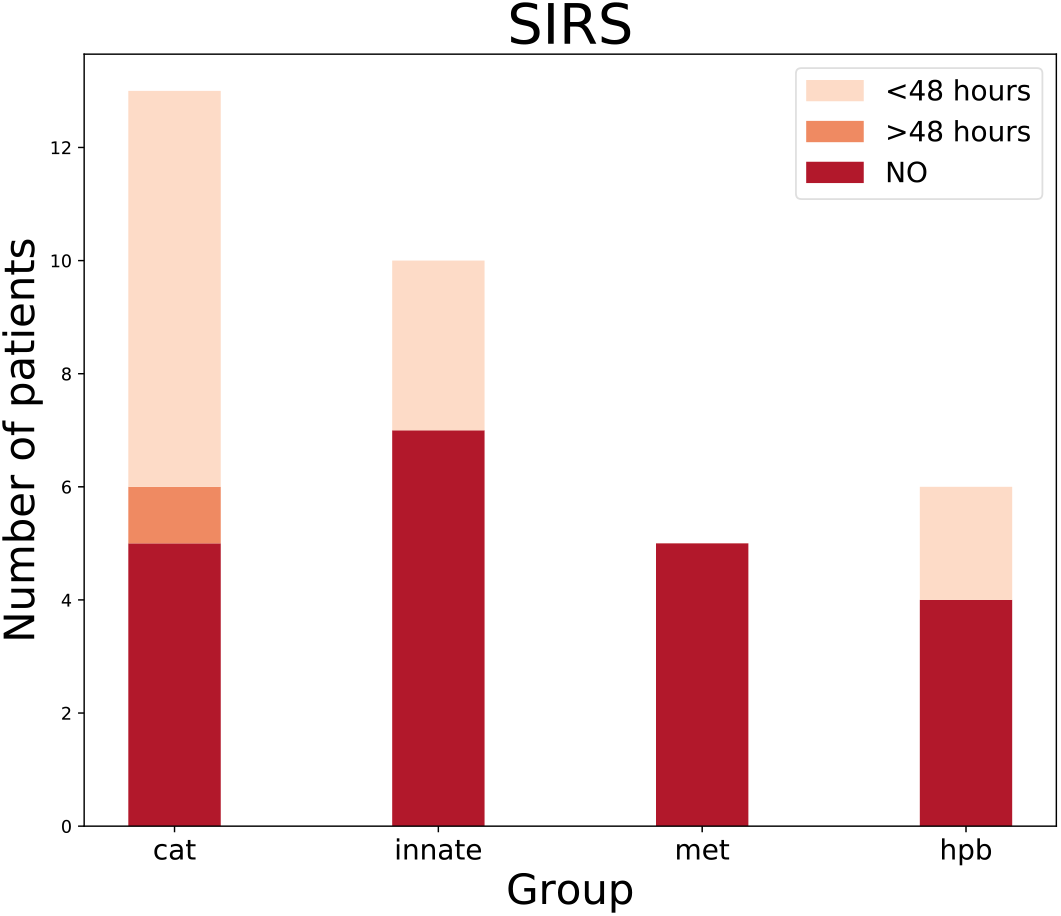
SIRS distribution per endotype. For each endotype, the number of patients is shown.

**Supplementary Figure 6.**
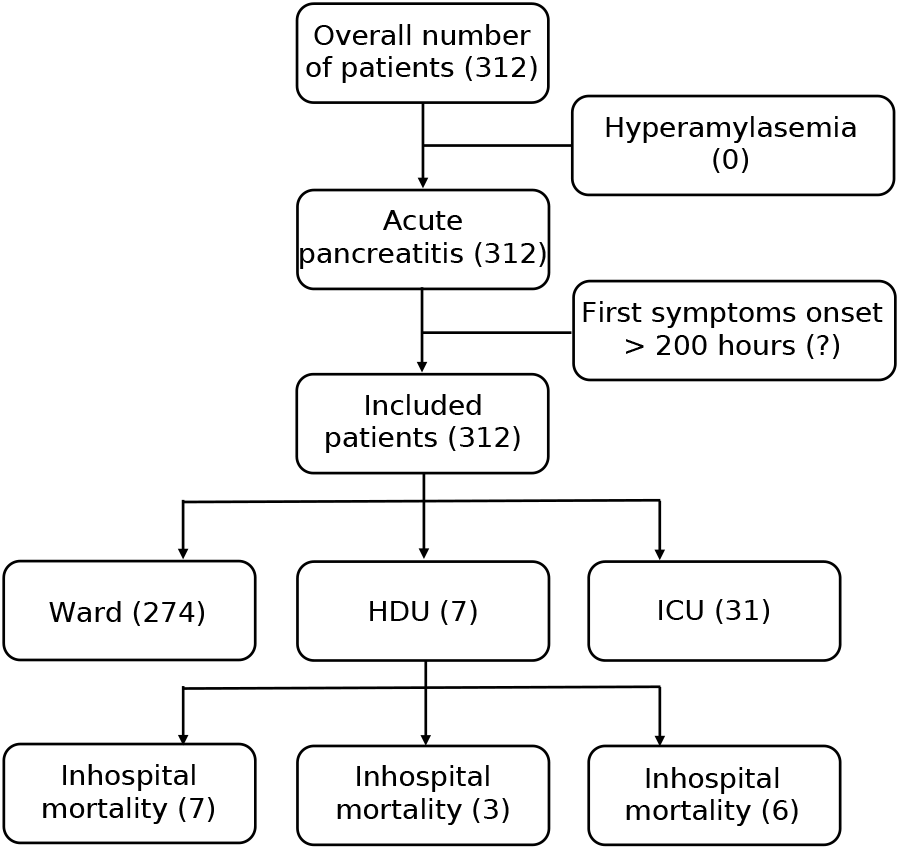
Flow diagram for the KAPVAL cohort. Study flow chart for patient samples and data from the KAPVAL study included in the analysis showing cohort structure.

**Supplementary Figure 7.**
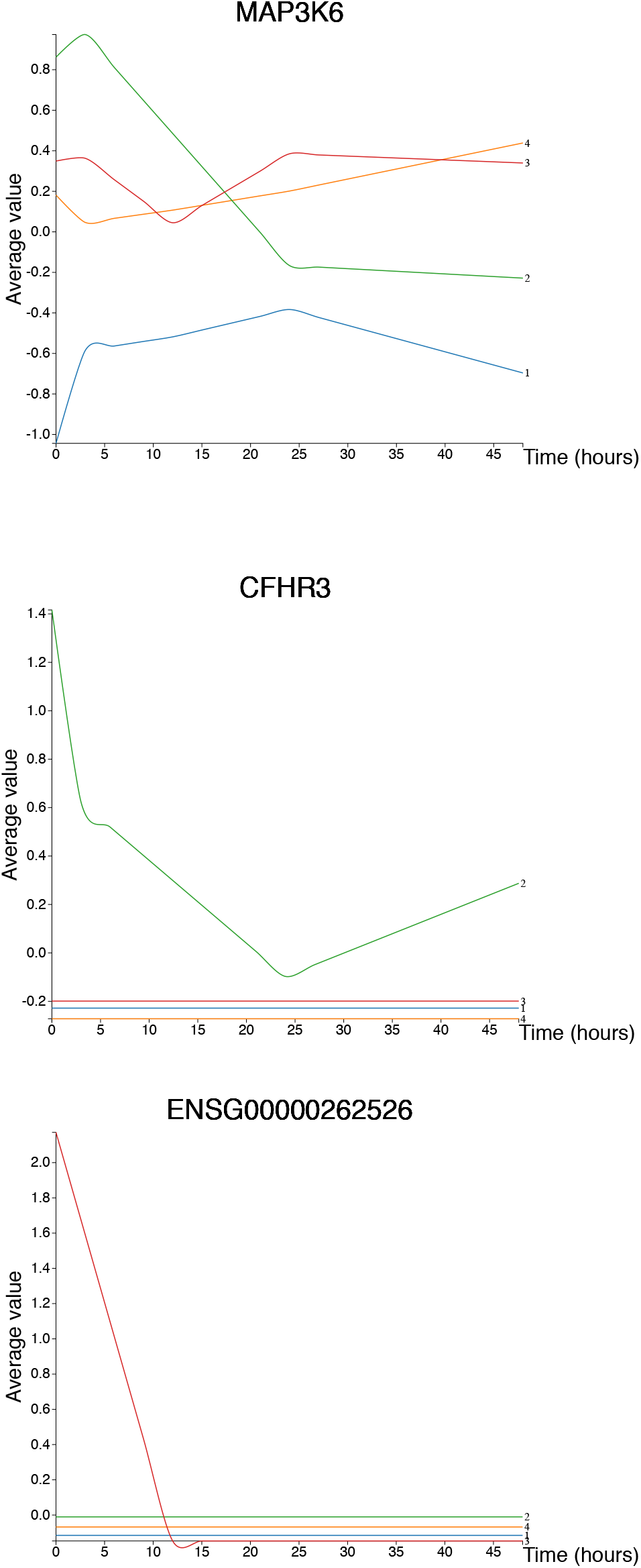
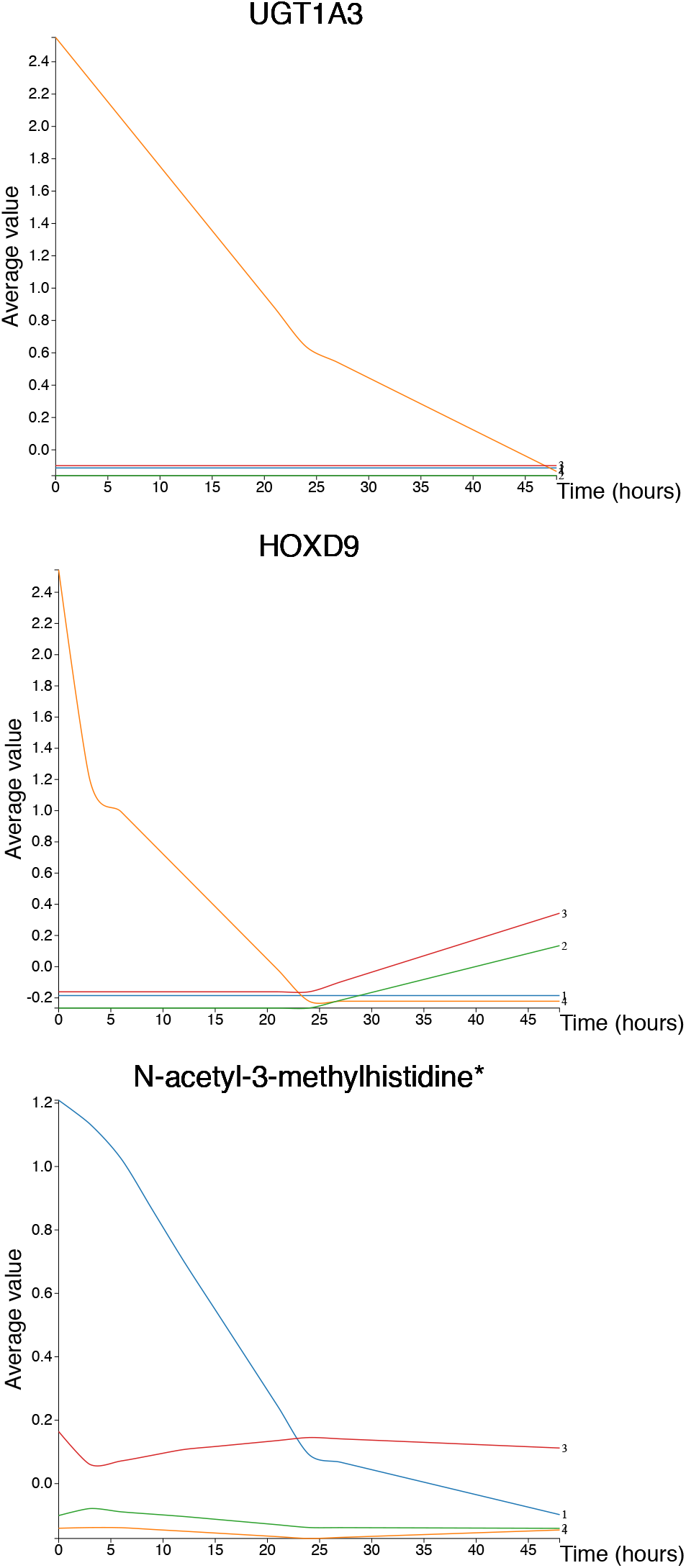
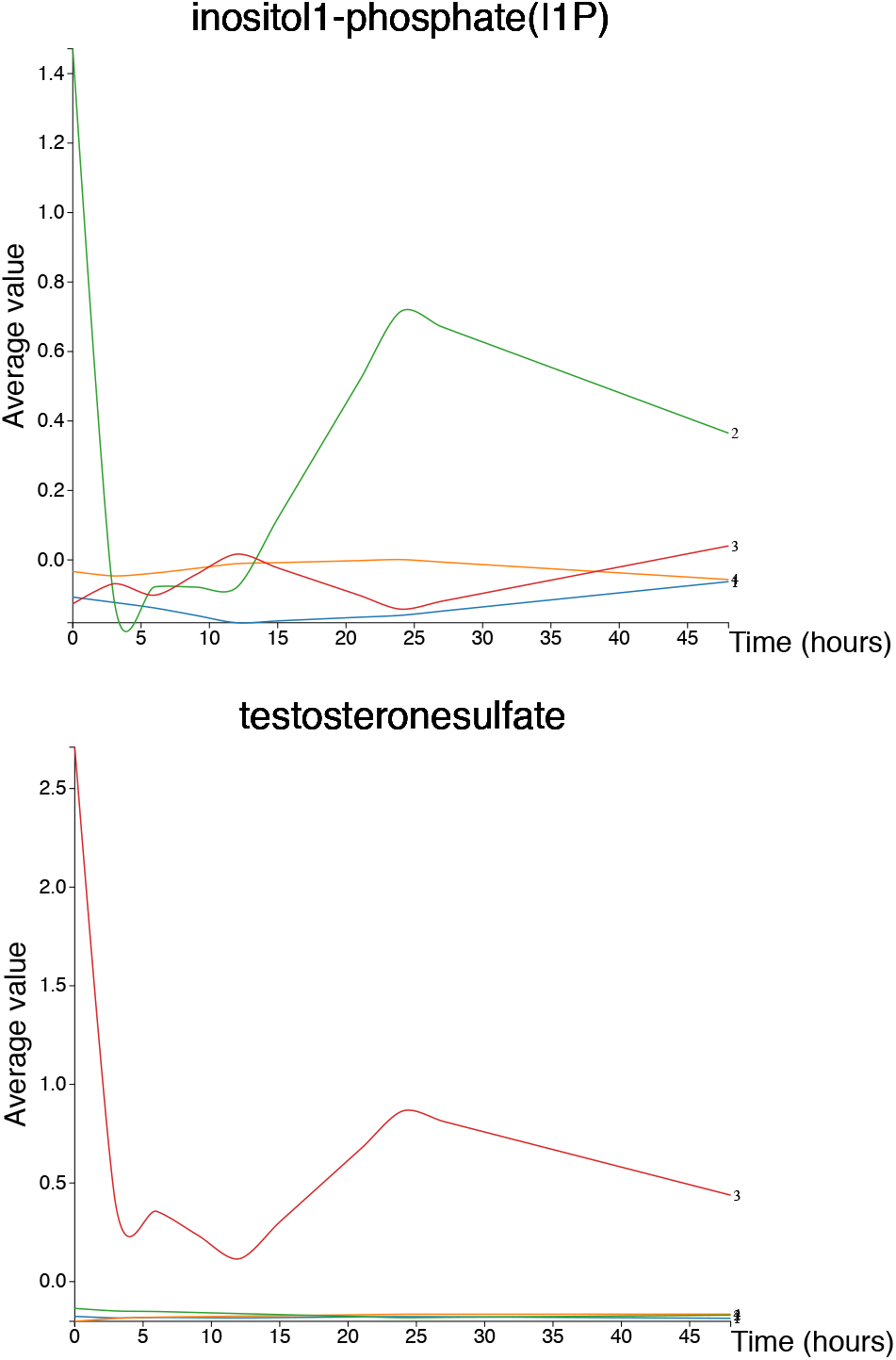
Time profiles per group for selected variables. Time series generated from average z-score value per time point per group identified using our AUC-PCA strategy. Top 2 variables per group selected from PLS-DA results (graphs produced using http://baillielab.net/pancreatitis/, username: pancreas and password: review).

**Supplementary Figure 8.**
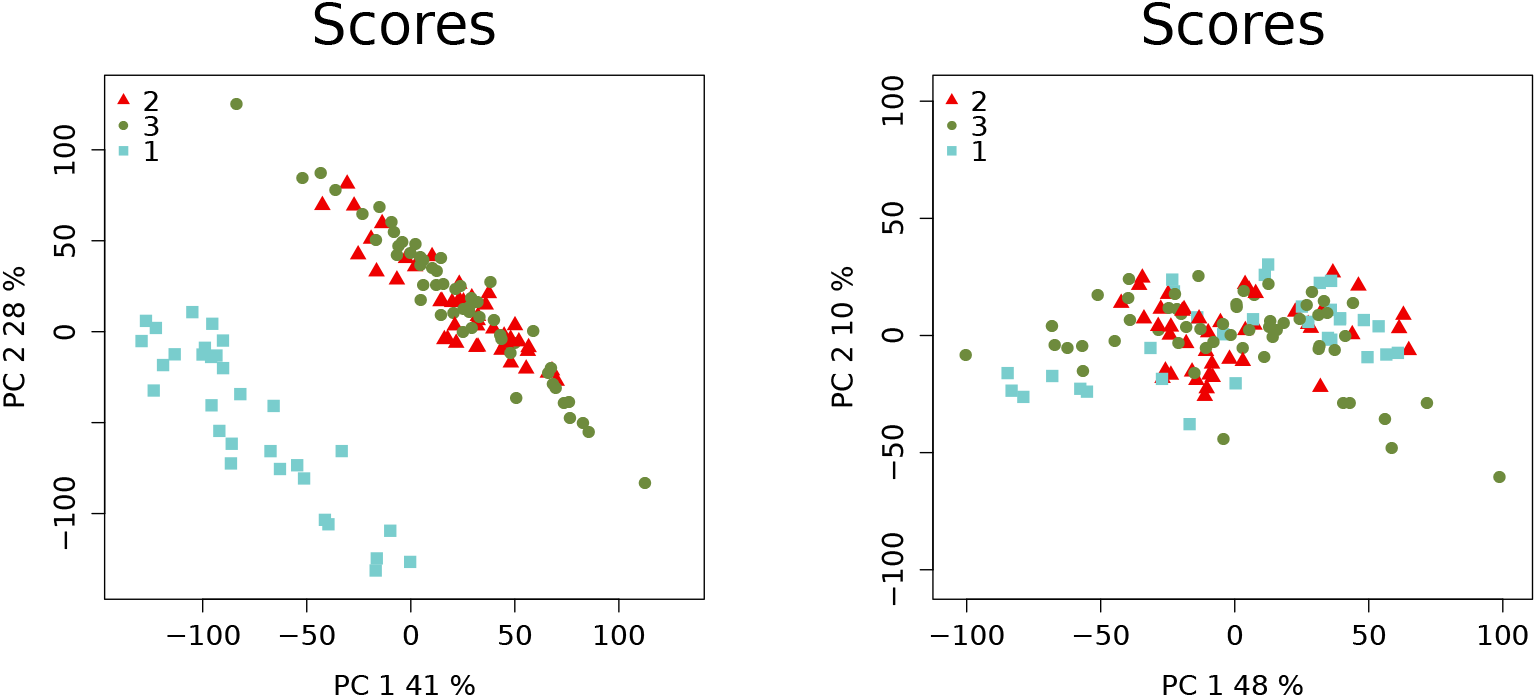
Batch effect correction for RNA-Seq data. PCA plots before and after batch effect removal. RNA-Seq counts values obtained using featureCounts, for coding genes only, are represented in the left figure. The same counts, after batch effect correction and FPKM normalisation are represented in the right figure.

**Supplementary Figure 9.**
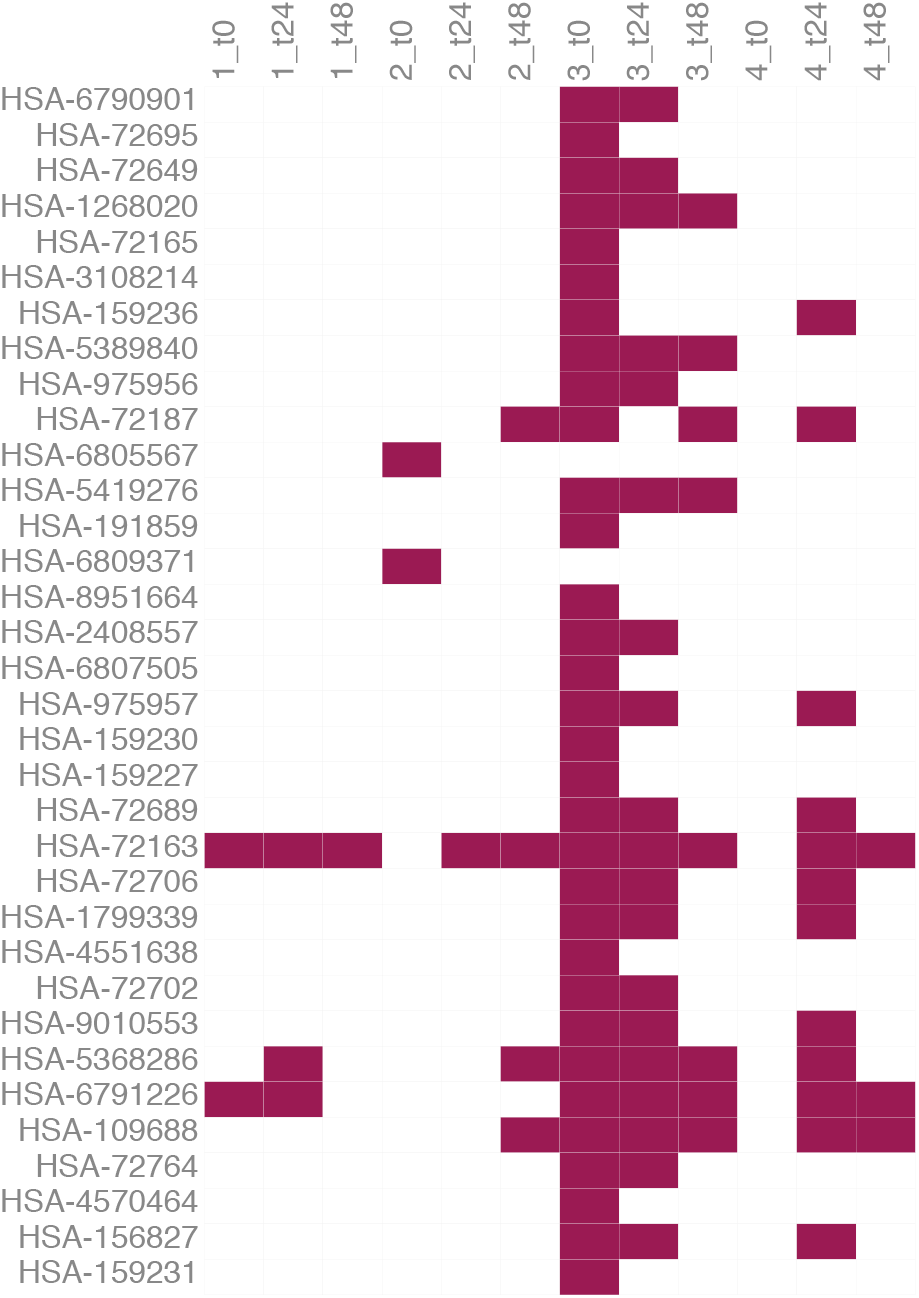
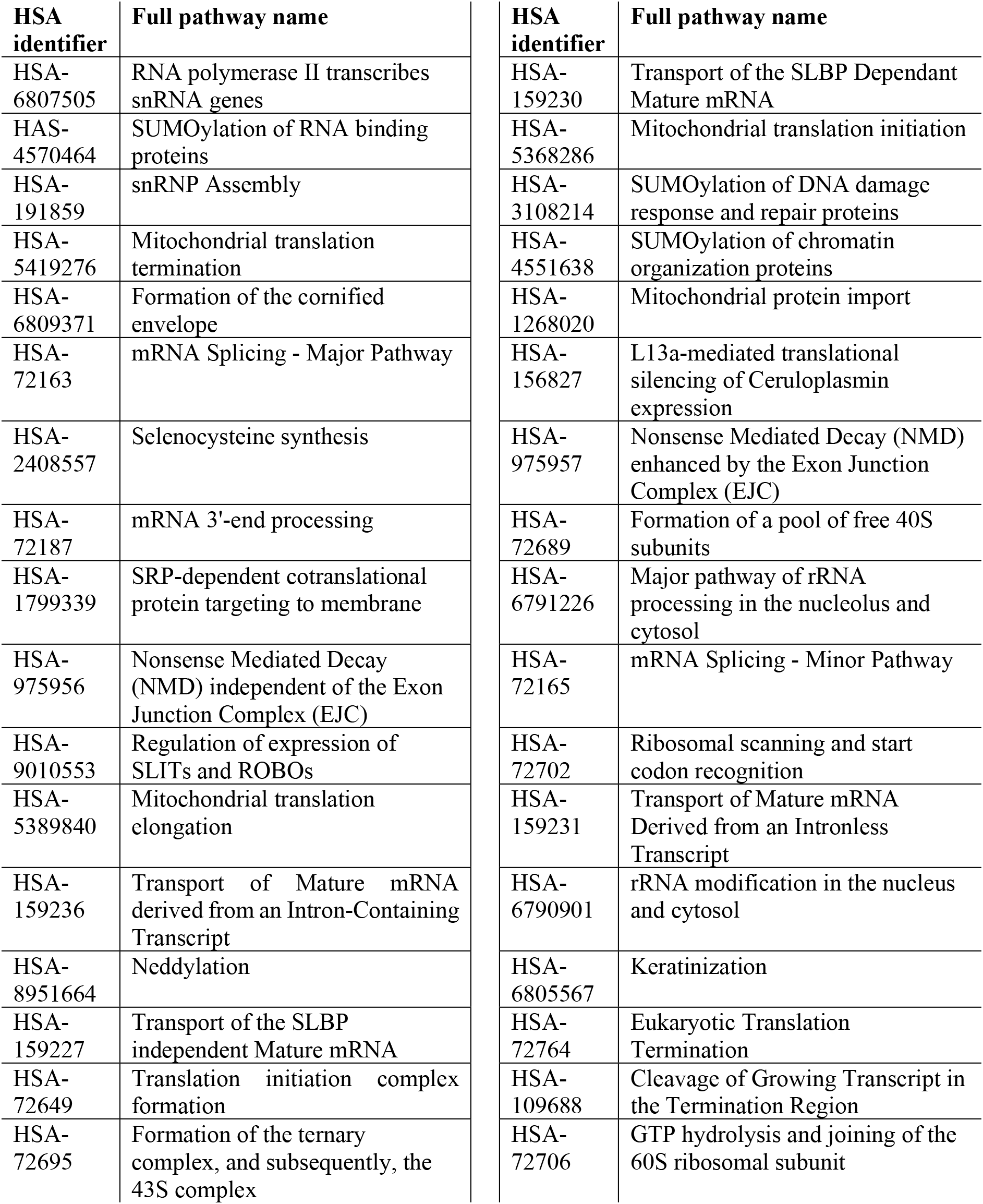
Enrichment results. Significant pathway terms (adjusted p-value threshold 0.01%) from enrichment results for each identified group based on variables lists selected using VIP scores. Pathway data extracted from Reactome database. Time point 0 was selected for assessment. Results for time points 24 and 48 hours are reported as well. Input variables were selected based on PLS-DA models and applying a threshold of 1 on VIP score values.

**Supplementary Figure 10.**
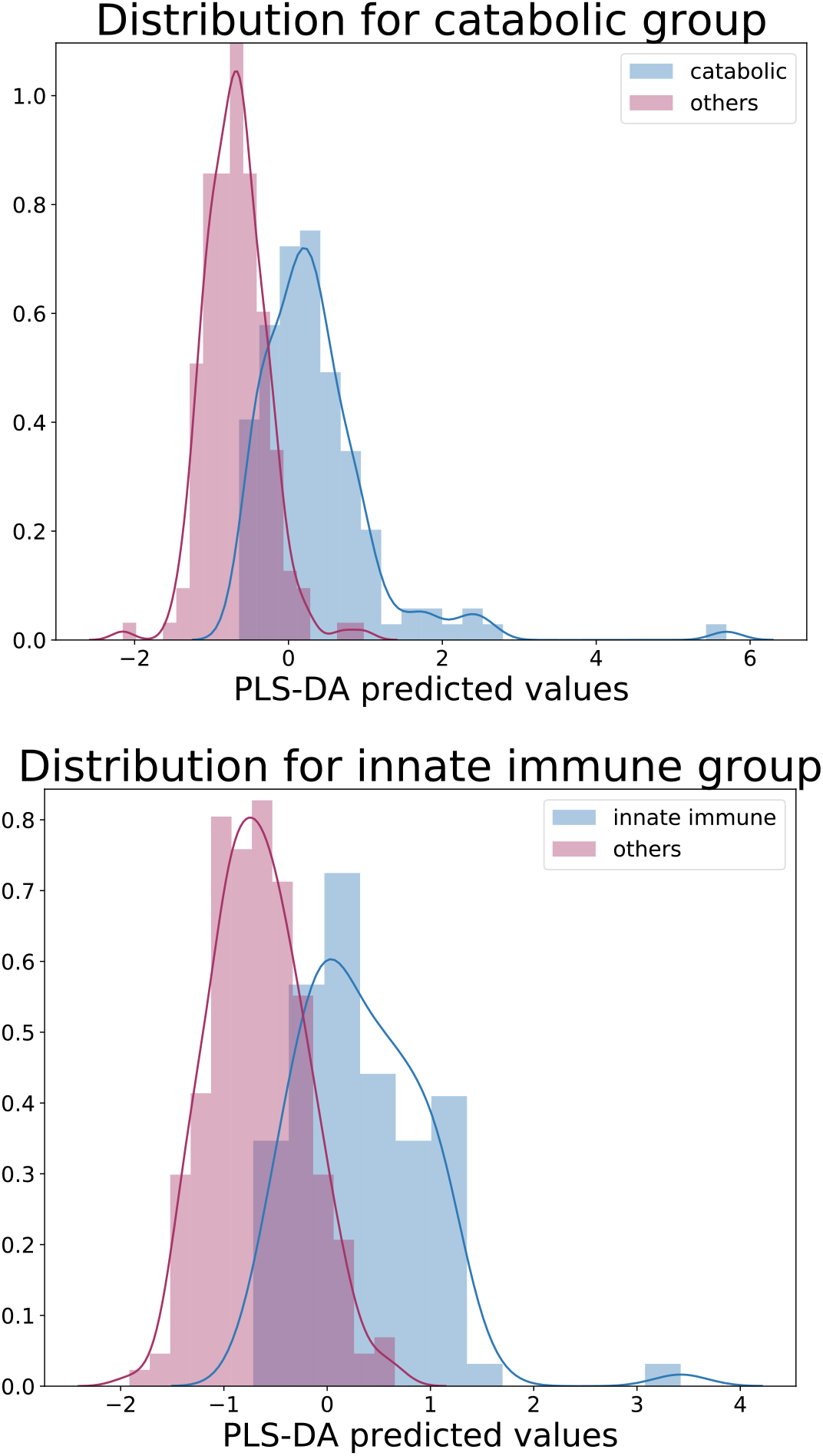
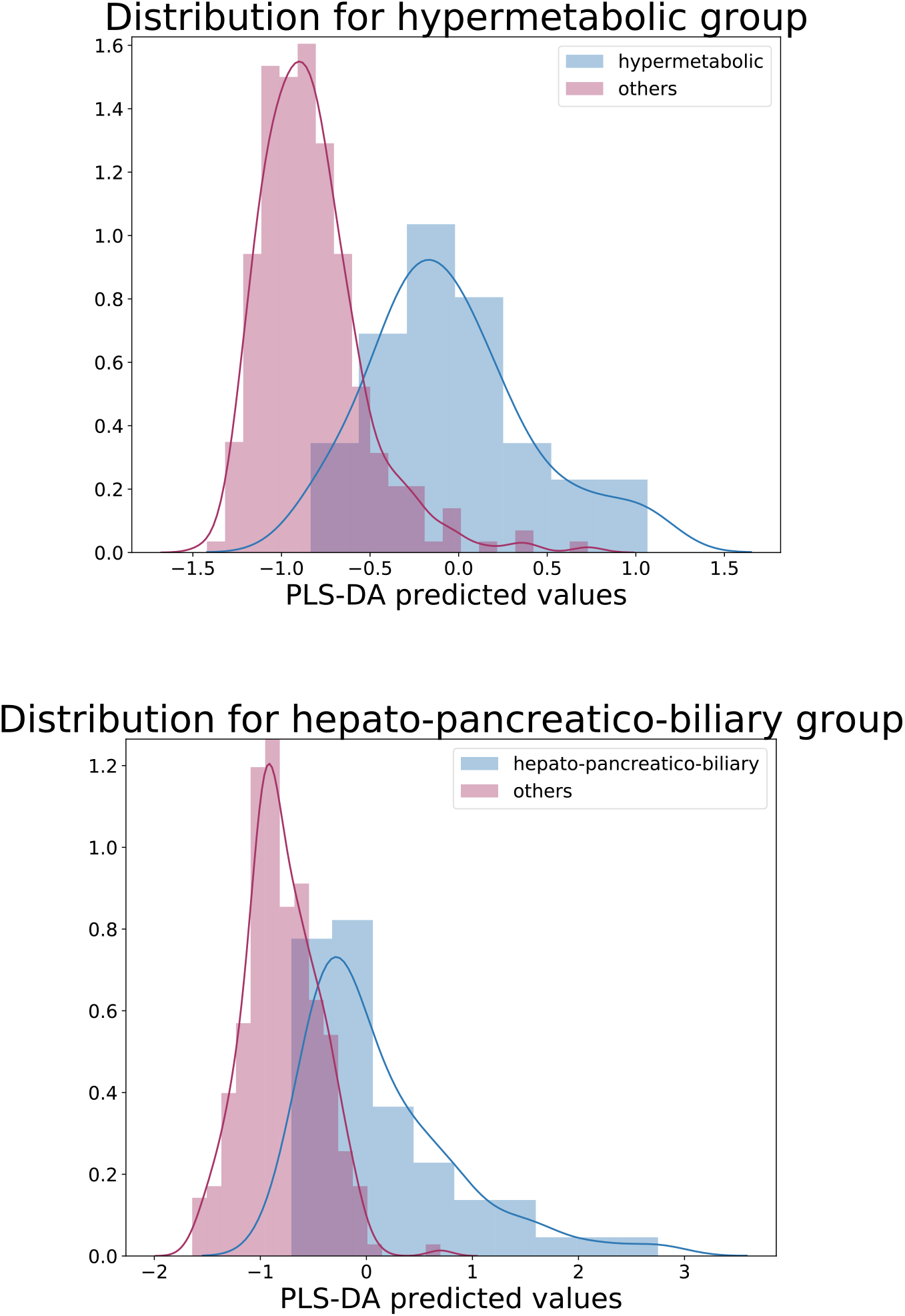
PLS-DA predicted values for each identified endotype. Distribution of PLS-DA predicted values for each identified endotype. For each endotype, distribution of predicted values for assigned and unassigned KAPVAL individuals are represented.

**Supplementary Figure 11.**
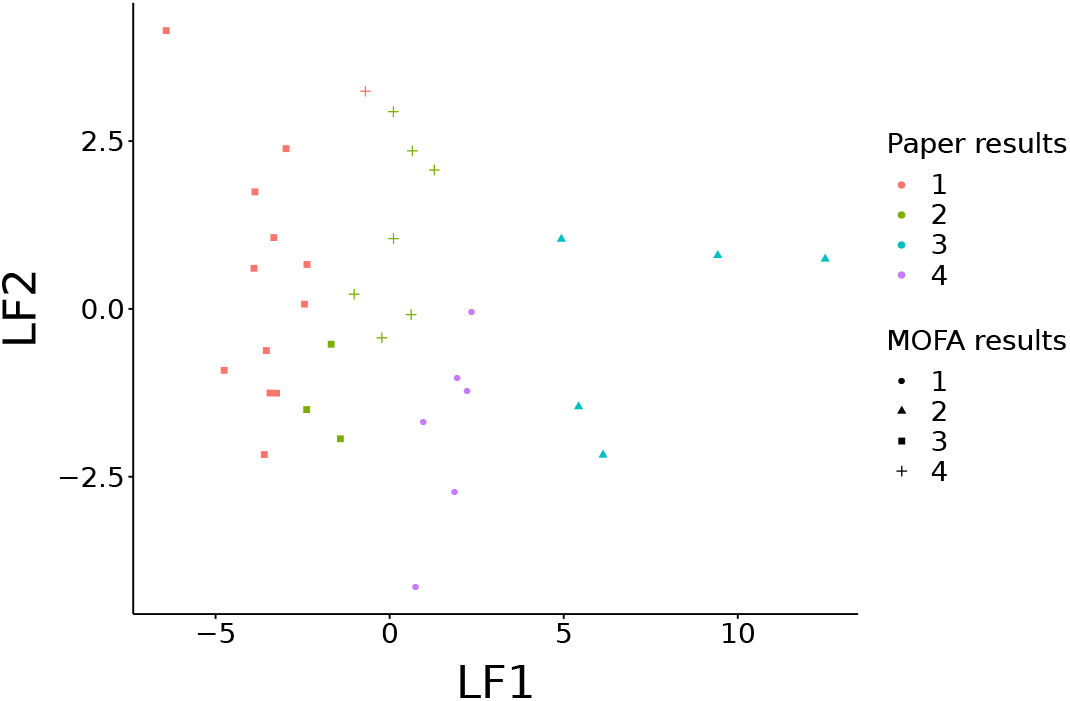
MOFA tools results compared to highlighted clusters with IMOFAP cohort data. Comparison of MOFAtools results with obtained clusters. AUC values used as input and a 4-cluster solution extracted from MOFA results using the first two latent features. Shapes are representative of clusters described in this paper and colours of MOFAtools predicted allocations.

**Supplementary Figure 12.**
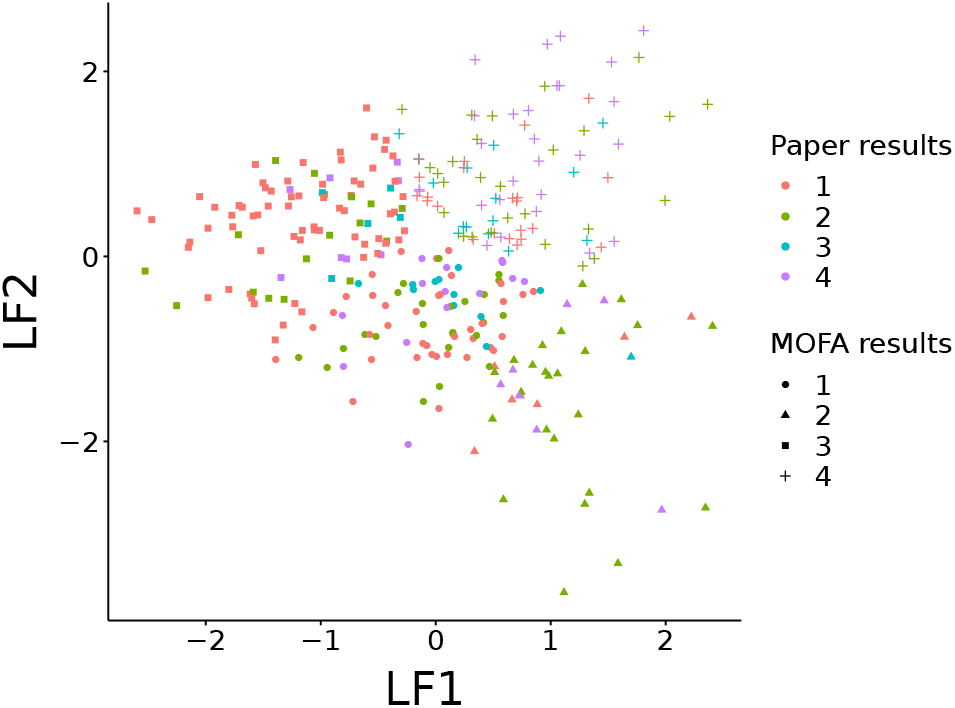
MOFA tools results compared to highlighted clusters with KAPVAL cohort data. Using KAPVAL metabolite data and selecting a 4-cluster solution, comparison of results. Shapes indicate results obtained using PLS-DA models and colours show MOFAtools results.

## Supplementary Tables

**Supplementary Table 1.**
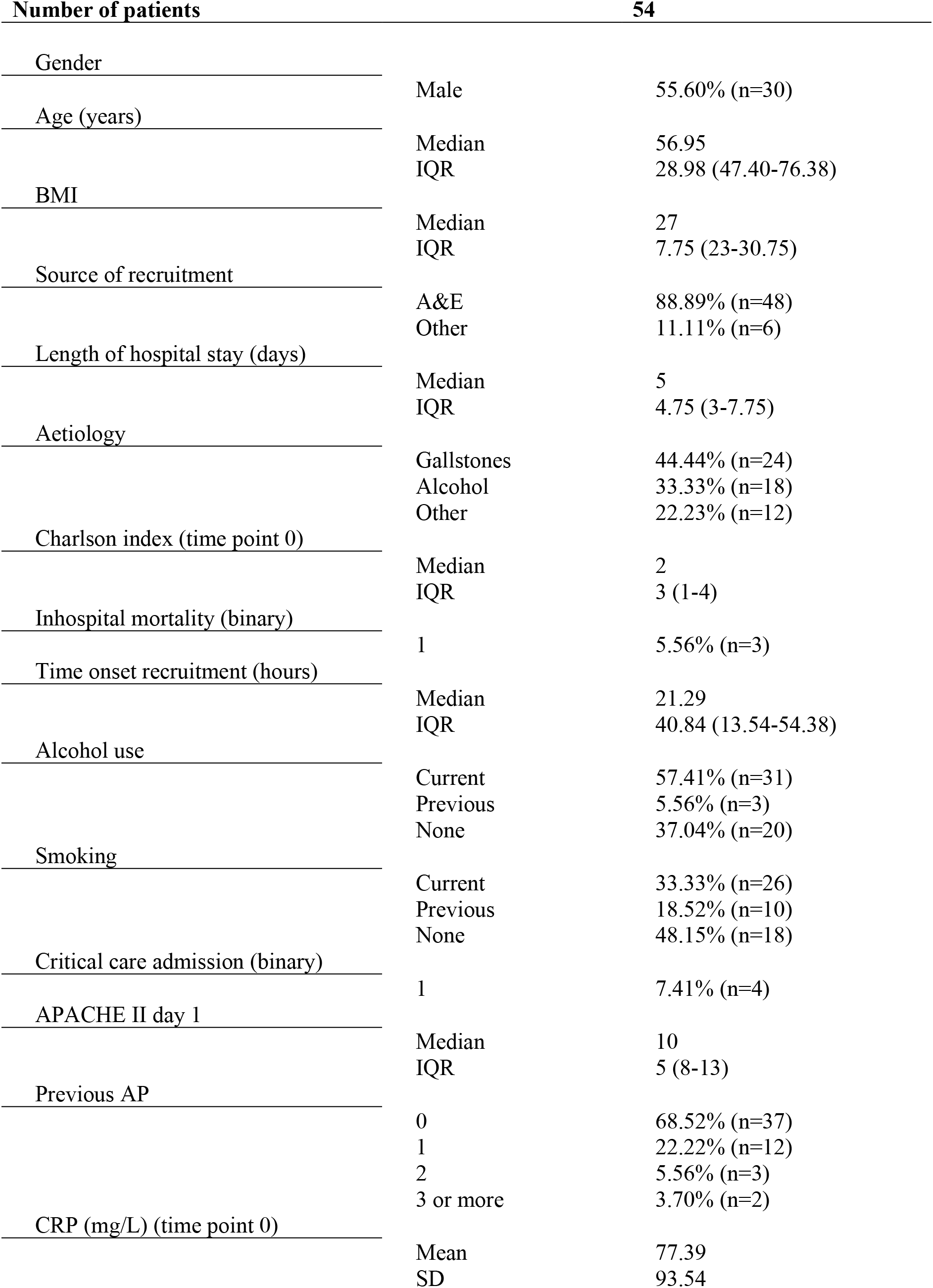
Demographics. Summary clinical data for included participants of the IMOFAP cohort. The cohort is fully described in Skouras et al (for n=57 AP patients).

**Supplementary Table 2.** Clustering similarities for 3-cluster solutions. For each time-series method, based on the 3-cluster solutions, comparisons were ran using Jaccard index values. Reported values are averaged Jaccard indexes. Values are displayed for each pairwise comparison.

| **Average Jaccard index** | AUC+PCA | PCA+Trajectory | Dynamic time warping |
| --- | --- | --- | --- |
| AUC+PCA | / | / | / |
| PCA+Trajectory | 0.31 | / | / |
| Dynamic time warping | 0.63 | 0.27 | / |

**Supplementary Table 3.** Clustering similarities for 5-cluster solutions. For each time-series method, based on the 5-cluster solutions, comparisons were ran using Jaccard index values. Reported values are averaged Jaccard indexes. Values are displayed for each pairwise comparison.

| **Average Jaccard index** | AUC+PCA | PCA+Trajectory | Dynamic time warping |
| --- | --- | --- | --- |
| AUC+PCA | / | / | / |
| PCA+Trajectory | 0.24 | / | / |
| Dynamic time warping | 0.46 | 0.21 | / |

**Supplementary Table 4.** Top pathways. Using global test, top 20 pathways (using KEGG data for gene, protein and metabolite data and FANTOM5 data for gene and protein data) for the AUC combined with PCA method. P-values obtained are reported along with pathway names/identifiers. Input data corresponds to time point 0.

| Pathway | p-value |
| --- | --- |
| hsa00190 Oxidative phosphorylation | 0 |
| hsa00230 Purine metabolism | 0 |
| hsa00240 Pyrimidine metabolism | 0 |
| hsa00510 N-Glycan biosynthesis | 0 |
| hsa00970 Aminoacyl-tRNA biosynthesis | 0 |
| hsa03008 Ribosome biogenesis in eukaryotes | 0 |
| hsa03010 Ribosome | 0 |
| hsa03013 RNA transport | 0 |
| hsa03015 mRNA surveillance pathway | 0 |
| hsa03018 RNA degradation | 0 |
| hsa03040 Spliceosome | 0 |
| hsa04010 MAPK signaling pathway | 0 |
| hsa04110 Cell cycle | 0 |
| hsa04120 Ubiquitin mediated proteolysis | 0 |
| hsa04141 Protein processing in endoplasmic reticulum | 0 |
| hsa04142 Lysosome | 0 |
| hsa04144 Endocytosis | 0 |
| hsa04146 Peroxisome | 0 |
| hsa04660 T cell receptor signaling pathway | 0 |
| hsa00280 Valine, leucine and isoleucine degradation | 5.79E-275 |

**Supplementary Table 5.** Heatmap compounds characteristics. Compounds detailed table for heatmap presented in figure 2b. As ordered in figure. Complete gene names were fetched using GeneCards resource and additional information using online resources as described in the main text.

| Heatmap compound | Complete name | Additional information |
| --- | --- | --- |
| GNAI1 | G Protein Subunit Alpha I1 | N-acetyl transferase activity |
| SPTSSB | Serine Palmitoyltransferase Small Subunit B | Tricarboxylic acid cycle |
| Citrulline | / | Sphingolipid biosynthesis |
| Dopamine sulfate (2) | / | Gastrointestinal dopamine metabolism |
| Testosterone sulfate | / |  |
| 5-acetylamino-6-amino-3-methyluracil | / | Caffeine metabolism |
| 5-acetylamino-6-formylamino-3-methyluracil | / |  |
| GGT2 | Gamma-Glutamyltransferase 2 | γ-glutamyl transferase; Glutathione homeostasis |
| URGCP-MRPS24 | URGCP-MRPS24 Readthrough |  |
| ENSG00000262526 | / | Protein coding |
| OR5D16 | Olfactory Receptor Family 5 Subfamily D Member 16 |  |
| CTAG1A | Cancer/Testis Antigen 1A |  |
| MYADML2 | Myeloid Associated Differentiation Marker Like 2 |  |
| Ribose | / |  |
| CELA2A | Chymotrypsin Like Elastase Family Member 2A | Pancreatic elastase-2 |
| HOXD9 | Homeobox D9 |  |
| OR6C6 | Olfactory Receptor Family 6 Subfamily C Member 6 |  |
| UGT1A3 | UDP Glucuronosyltransferase Family 1 Member A3 | Associated with Gilbert-type hyperbilirubinemia |
| SLCO1B7 | Solute Carrier Organic Anion Transporter Family Member 1B7 (Putative) | Cysteine-type endopeptidase |
| USP17L18 | Ubiquitin Specific Peptidase 17-Like Family Member 18 | Liver-specific organic anion transporter; Bile secretion |
| DMRTC1 | DMRT Like Family C1 |  |
| CGB3 | Chorionic Gonadotropin Subunit Beta 3 |  |
| N-acetyl-1-methylhistidine* | / | Amino acid metabolism; Rhabdomyolysis; Renal failure |
| N-acetyl-3-methylhistidine* | / |  |
| PPP1R42 | Protein Phosphatase 1 Regulatory Subunit 42 |  |
| SLC16A8 | Solute Carrier Family 16 Member 8 | Lactate transporter; Ketone body transporter |
| KRTAP6-3 | Keratin Associated Protein 6-3 | Muscle-specific actin binding protein upregulated during muscle injury |
| XIRP1 | Xin Actin Binding Repeat Containing 1 |  |
| MAP3K6 | Mitogen-Activated Protein Kinase Kinase Kinase 6 | Apoptosis signaling |
| BICDL2 | BICD Family Like Cargo Adaptor 2 |  |
| ZBBX | Zinc Finger B-Box Domain Containing |  |
| CFHR3 | Complement Factor H Related 3 | Heparin-binding; Complement regulation |
| Inositol 1-phosphate (I1P) | / | Inositol biosynthesis |
| HOXD3 | Homeobox D3 | Increases immune cell adherence; Overexpression upregulates glycoprotein IIb/IIIa |
| SPEM1 | Spermatid Maturation 1 |  |
| C6orf15 | Chromosome 6 Open Reading Frame 15 | Putative heparin/fibronectin binding |
| TRIM48 | Tripartite Motif Containing 48 | Interferon-γ signalling (oxidative stress/apoptosis signal-reducing kinase 1) |
| REG3A | Regenerating Family Member 3 Alpha | Bactericidal C-type lectin; Known as pancreatitis-associated protein |
| PPP1R3A | Protein Phosphatase 1 Regulatory Subunit 3A | Genetic association with type 2 DM and familial partial lipodystrophy 3 |

**Supplementary Table 6.** Demographics. Summary clinical data for included participants of the KAPVAL cohort.

| Number of patients |  | 312 |
| --- | --- | --- |
| Gender | Male | 46.79% (n=146) |
| Age (years) | Median | 56.00 |
|  | IQR | 30.25 (40.75-71.00) |
| Inhospital mortality (binary) | 1 | 5.13% (n=16) |
| Critical care admission (binary) | 1 | 12.18% (n=38) |
| CRP (mg/L) | Mean | 47.62 |
|  | SD | 79.85 |

## Notes

#### Summary of Updates

Updated manuscript after peer review

